# Chromatin organization in early land plants reveals an ancestral association between H3K27me3, transposons, and constitutive heterochromatin

**DOI:** 10.1101/827881

**Authors:** Sean A. Montgomery, Yasuhiro Tanizawa, Bence Galik, Nan Wang, Tasuku Ito, Takako Mochizuki, Svetlana Akimcheva, John Bowman, Valérie Cognat, Laurence Drouard, Heinz Ekker, Syuan-Fei Houng, Takayuki Kohchi, Shih-Shun Lin, Li-Yu Daisy Liu, Yasukazu Nakamura, Lia R. Valeeva, Eugene V. Shakirov, Dorothy E. Shippen, Wei-Lun Wei, Masaru Yagura, Shohei Yamaoka, Katsuyuki T. Yamato, Chang Liu, Frédéric Berger

**Author notes:** SAM and YT contributing equally and share first authorship.

## Abstract

Genome packaging by nucleosomes is a hallmark of eukaryotes. Histones and the pathways that deposit, remove, and read histone modifications are deeply conserved. Yet, we lack information regarding chromatin landscapes in extant representatives of ancestors of the main groups of eukaryotes and our knowledge of the evolution of chromatin related processes is limited. We used the bryophyte *Marchantia polymorpha*, which diverged from vascular plants 400 Mya, to obtain a whole chromosome genome assembly and explore the chromatin landscape and three-dimensional organization of the genome of early land plants. Based on genomic profiles of ten chromatin marks, we conclude that the relationship between active marks and gene expression is conserved across land plants. In contrast, we observed distinctive features of transposons and repeats in *Marchantia* compared with flowering plants. Silenced transposons and repeats did not accumulate around centromeres, and a significant proportion of transposons were marked by H3K27me3, which is otherwise dedicated to the transcriptional repression of protein coding genes in flowering plants. Chromatin compartmentalization analyses of Hi-C data revealed that chromatin regions belonging to repressed heterochromatin were densely decorated with H3K27me3 but not H3K9 or DNA methylation as reported in flowering plants. We conclude that in early plants, H3K27me3 played an essential role in heterochromatin function, suggesting an ancestral role of this mark in transposon silencing.

## INTRODUCTION

In eukaryotes, the evolution of histones that assemble with DNA into nucleosomes generated chromatin with a more diverse composition and complex organization compared to that found in prokaryotes [1, 2]. Post-translational modifications of core histones that form nucleosomes contribute to the complexity and flexibility of chromatin [3]. The characterization of such modifications, marking transcriptionally active and inactive regions of the genome, has furthered insights into the functional organization of eukaryotic chromatin. In flowering plants, extensive meta analyses of histone modifications profiles in *Arabidopsis thaliana* highlighted the association of H3K4me3, H3K36me3, and H3 acetylation with gene expression, while H3K27me3 marks transcriptional repression and H3K9 methylation is associated with DNA methylation (5’methyl Cytosine) marking silenced transposons [4].

The three-dimensional (3D) organization of domains where distant regions of chromatin connect is revealed by genomic methods such as Hi-C [5] and genome architecture mapping [6]. The 3D organization of the genome of flowering plants analyzed by classical cytological methods and Hi-C showed a wide variety of nuclear organization patterns [7, 8]. The diversity in chromatin organization suggests that during evolution, genome organization changed and diversified depending on genome duplications, size, and relative content of transposons versus genes. It is therefore important to extend investigations of 3D genome organization to a larger number of species representative of extant ancestral lineages to understand how genome architecture evolved in eukaryotes.

Bryophytes, comprised of liverworts, mosses, and hornworts, represent ancient lineages of land plants which diverged from the vascular plant lineage over 400 Mya [9]. Analysis of the genome sequences of the liverwort *Marchantia polymorpha* and the moss *Physcomitrella patens* showed that genes encoding pathways related to histone modifications are broadly conserved in land plants [10], but that heterochromatic islands of transposons and repeats alternate with genes without a clear demarcation of a region enriched in transposons around centromeres [11]. This contrasts with the vast accumulation of transposons and repeats around centromeres described in *Arabidopsis* and many species of flowering plants [12, 13]. Yet, the lack of Hi-C maps and the limited knowledge of chromatin modifications profiles in bryophytes has limited our understanding of the ancestral functional organization of chromatin in land plants.

We obtained a new full chromosome assembly of the genome of the liverwort *Marchantia polymorpha* (male accession Tak-1) with an update of annotations, which will be publicly accessible as reference genome version 5.1 for this species. Here, we present a new set of extensive profiles of key chromatin marks as well as 3D chromatin organization patterns obtained by Hi-C. Altogether, our observations lead to a model of chromatin organization in early land plants, revealing that considerable changes arose during the evolution of vascular plants.

## RESULTS

### A chromosome assembly of the *Marchantia* genome

The previous version of the nuclear genome of *Marchantia polymorpha* (version 3.1) comprised 2,957 scaffolds with 19,138 nuclear encoded protein-coding genes [10]. We obtained a new set of scaffolds of the genome from the male accession Tak-1 using long-read sequencing and assembled them at a chromosomal scale using Hi-C (Figure S1). Overall, this newly assembled Tak-1 genome, referred to as *Marchantia polymorpha* version 5.1, contains 218.7 Mb, including 215.8 Mb jointly covered by the autosomes and sex chromosome (chromosome V), and can be accessed at MarpolBase (http://marchantia.info/). A total of 200 Mb genomic regions showed high sequence identity (>99% identity) against the version 3.1 genome. The majority of the additional 17.7 Mb was accounted for by repetitive regions (14 Mb), while the remaining 3.7 Mb showed lower similarity or no homology against the version 3.1 genome. Markers associated in distinct genetic linkage groups were identified between the two accessions Tak-1 and Tak-2 (Table S2). The linkage groups and linear order of the vast majority of these genetic markers were fitted correctly with the chromosomes assembled in version 5.1 (Table S2). This genetic map at low resolution validated the overall structure of the physical whole chromosome genome assembly.

The version 5.1 genome harbors 19,421 predicted protein-coding loci with 24,751 transcript models including isoforms (Table S1). Among them, 24,078 transcript models were carried over from the version 3.1 genome, and 673 were newly identified by *de novo* prediction and manual inspection. We also curated new 303 transcript models based on expression evidence from RNA-seq and Iso-seq. The completeness of the gene set was assessed using BUSCO [14], estimating that 97.6% (296) out of 303 universal single-copy orthologs for eukaryotes were present, the same level as the version 3.1 genome. We adopted a new series of unique gene identifiers following the guidelines established for the *Arabidopsis* genome. Examples of newly identified genes include gene clusters such as the NNP family, nitrate/nitrite transporters (Mp5g10710, Mp5g10760, Mp5g10780, Mp5g10790), metalloproteases (Mp8g14490, Mp8g14520, Mp8g14560, Mp8g14610), and DEAD-box family RNA helicases (Mp4g13200, Mp4g13270, Mp4g13330). These regions were missing or fragmented into different scaffolds in the version 3.1 genome, indicating the advantage of the version 5.1 assembly leveraged by long-read sequencing in reconstructing such repetitive regions. We also identified comprehensive lists tRNAs, miRNAs, transposons, and repeats (Table S2).

The male-specific sex chromosome V of *Marchantia* consists of two parts, each of which has distinctive sequence content, YR1 and YR2 [15]. YR1 is highly enriched in repeats unique to chromosome V [15, 16]. Version 5.1 includes two novel regions of the V chromosome, a 506-kb region between Contig-A and Contig-B, and a 1.3-Mb region at the distal end of Contig-A from Contig-B. The 1.3-Mb region contains blocks of the V-specific repeats (Figure S2), most likely representing part of YR1. The extremely high repeat content still prevented this region from being fully assembled and reconstructed. Interestingly, copies of rDNA were found among the blocks of the V-specific repeats (Figure S2). Two types of rDNA were reported to be present in the *Marchantia* genome, one autosomal and the other U-chromosomal [17]. The V-chromosomal copies were more similar to the autosomal (99.64%) than to the U-chromosomal copies (97.02%). Unlike the autosomal and U-chromosomal rDNAs, the V-chromosomal rDNAs do not form a regular tandem array suggesting potential for distinct epigenetic regulation as shown for distinct rDNA clusters in *Arabidopsis* [18].

### Telomeres, centromeres, and overall nuclear organization

Telomeres of *Marchantia polymorpha* are composed of tandem arrays of TTTAGGG repeats similar to that identified in *Marchantia palaeceae* [19]. To gauge the size of telomere tracts, we performed terminal restriction fragment analysis and observed that *Marchantia* telomeres are longer than in *Physcomitrella* and shorter than in *Arabidopsis* (Figure S3A). We concluded that *Marchantia* telomeres are comparable with those of most other plants [19–21].

In most flowering plants, centromeres are comprised of specific satellite repeats interspersed with transposons and surrounded by a pericentromeric region enriched in transposons. We identified centromeric repeats composed of 162 bp satellite DNA (Figure S3B). This size is within the range found in other land plants [22] and compatible with the typical shorter length of DNA associated with centromeric nucleosomes [23]. These repeats were found close to the center of each autosome (Figure S3C). The presence of a potential CENP-B box in the repeat (Figure S3B) strengthens the similarity of this repeat to other identified centromeric repeats [22]. Beyond the satellite repeats, long terminal repeats (LTR) retrotransposons accumulate in centromeres and pericentromeres of flowering plants and animals [24–26]. In contrast, in *Marchantia* we did not find LTR transposons in proximity of the centromeres but only the specific family LINE/RTE-X, which showed a sharp peak, surrounding centromeres of each chromosome (Figure S3C). These data indicate that *Marchantia* has monocentric centromeres marked by short repeats as described in the majority of land plants, but the extent of these repeats and the lack of LTR transposons do not define an extended pericentric region as observed in many flowering plants.

With the knowledge of *Marchantia* centromeric and telomeric regions, we designed probes to examine their distribution in interphase nuclei in the vegetative thallus. We found up to nine dots marked by the centromeric repeat probes, which showed a dispersed localization (Figure 1A). Telomeres were located as eighteen dots situated at the end of each chromosome in metaphase (Figure 1B). In interphase, telomeres often clustered to form a single speckle (Figure 1C). A similar conformation, called “bouquet”, has been reported in meiotic maize, wheat, and rice cells [27–29]. However, in contrast to bouquet conformation described in flowering plants, the telomere gathering in *Marchantia* nuclei did not display a specific association of telomeres with the nuclear periphery (Figure 1C).

**Figure 1.**
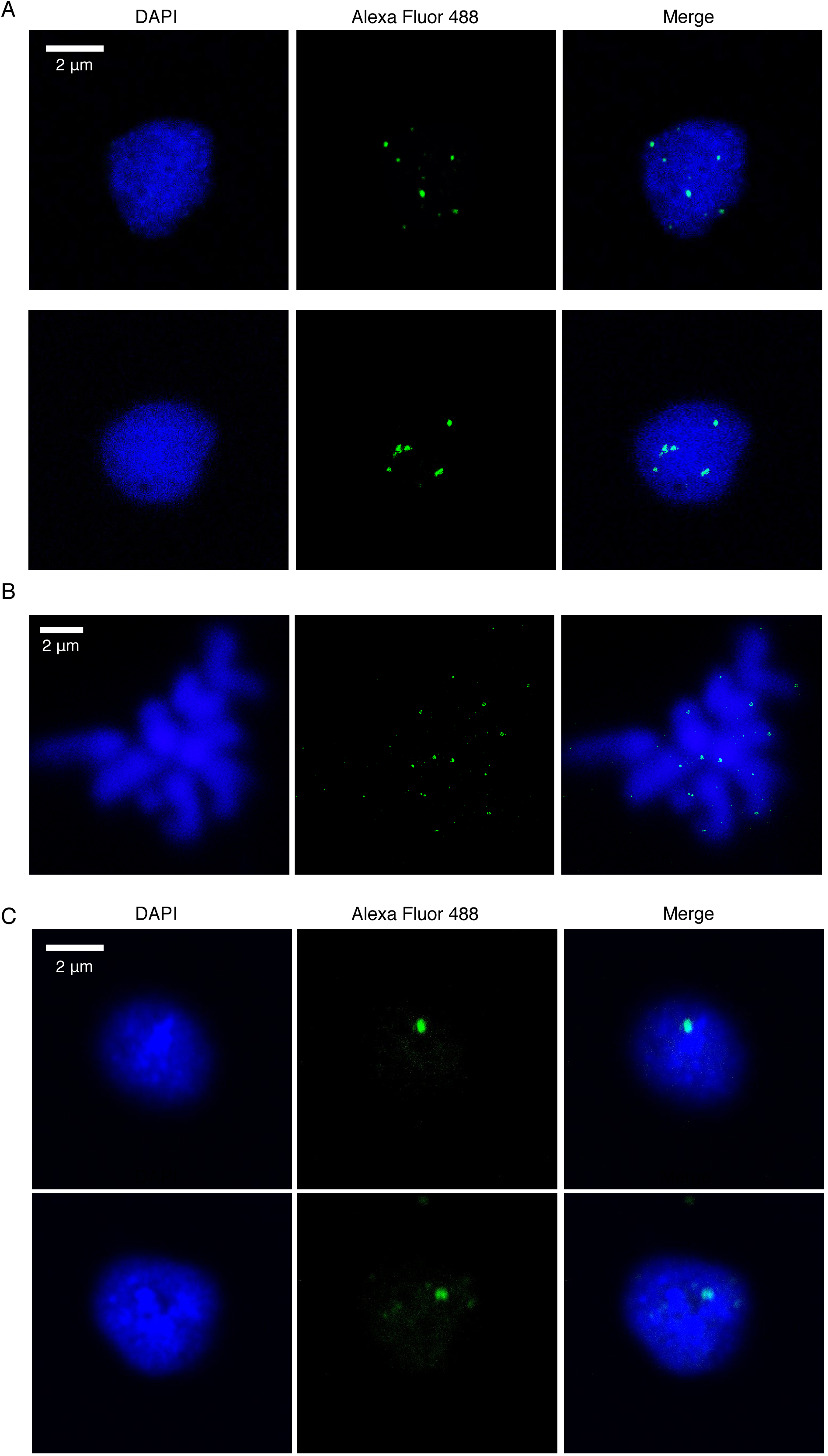
Distribution patterns of centromeric repeats and telomeres in *Marchantia*. (A) Distribution of centromeric repeats in Tak-1 nuclei isolated from vegetative thalli. Probes labeled with digoxigenin were hybridized with Tak-1 chromosome spread preparations and visualized with Alexa Fluor 488. (B) Confirmation of telomere probes’ specificity by using chromosome spread. (C) Distribution of telomeres in Tak-1 nuclei isolated from vegetative thalli. Probes labeled with digoxigenin were hybridized with Tak-1 chromosome spread preparations and visualized with Alexa Fluor 488.

To examine the spatial organization of euchromatin versus heterochromatin, we immunostained *Marchantia* and *Physcomitrella patens* nuclei with antibodies against histone modifications typical of constitutive heterochromatin (H3K9me1 and H3K27me1), facultative heterochromatin (H3K27me3), and euchromatin (H3K36me3 and H3K4me3) as defined in *Arabidopsis* [4]. The distribution of DNA in *Marchantia* is more punctate, with many small foci and several larger ones (Figure S4A), in comparison to the smooth and homogeneous distribution of DNA in *Physcomitrella patens* (Figure S4B). In *Marchantia* nuclei, heterochromatic regions, denoted by denser staining, tend to overlap with H3K9me1 and H3K27me1 but also surprisingly with H3K27me3. These heterochromatic regions do not form clear compact structures comparable to chromocenters described in *Arabidopsis* and other flowering plants. In *Physcomitrella* and to some degree in *Marchantia*, the euchromatic mark H3K36me3 tends to be excluded from heterochromatic regions and is remarkably enriched at the periphery of nuclei, while heterochromatic marks tend to be located at more central locations.

### Organization of chromatin profiles

Using CUT&RUN [30, 31] in *Marchantia polymorpha*, we obtained genomic profiles of eight histone modifications (H3K9me1, H3K27me1, H3K9ac, H3K14ac, H3K4me1, H3K36me3, H3K4me3, and H3K27me3), one histone variant (H2A.Z), and H3. This set of histone modifications together with data available for DNA methylation [32] and transcriptional activity [10], can be accessed at at MarpolBase (http://marchantia.info/). This comprehensive and integrated dataset enabled us to draw comparisons with chromatin states in *Arabidopsis* [4]. Biological replicates tended to cluster together in a Pearson correlation matrix (Figure S5A) and marks typically considered active (H3K9ac, H3K14ac, H3K36me3) or repressive (H3K9me1, H3K27me1) grouped amongst themselves (Figure S5B). Interestingly, H3K27me3 was quite distinct from other marks and correlated most strongly with H3K4me3 and H2A.Z. Accordingly, H3K27me3 peaks overlapped primarily with H3K4me3 and H2A.Z peaks (Figure S5C) but not with DNA methylation in CG, CHG, and CHH contexts [32], which were most strongly associated with H3K9me1 and H3K27me1 (Figure S5D).

Each of the chromatin profiles was spread evenly across chromosomes (Figures 2A and 2B) following the even distribution of transposons and genes. Peaks of H3K9me1 and H3K27me1 were enriched on ribosomal RNA coding genes, satellites, repeats, and transposons (Figures 2C and 2D). In flowering plants, centromeres are surrounded by heterochromatic pericentromeric regions marked by DNA methylation, H3K9me1, H3K9me2, and H3K27me1, that target multiple families of transposons [4, 13, 24, 33]. Such accumulation was not detected around centromeres in *Marchantia* (Figure 2A) and we concluded that there is no detectable pericentric heterochromatin in *Marchantia*. Strikingly, 60% of the peaks of H3K27me3 were found on repeats and transposons while the remaining peaks were associated with genes (Figure 2C). All other chromatin modifications profiled were primarily associated with genes with a notable enrichment of H3K36me3 over the coding sequence and 3’UTR while the 5’UTR is relatively more enriched in H3K9ac (Figure 2C and 2D).

**Figure 2.**
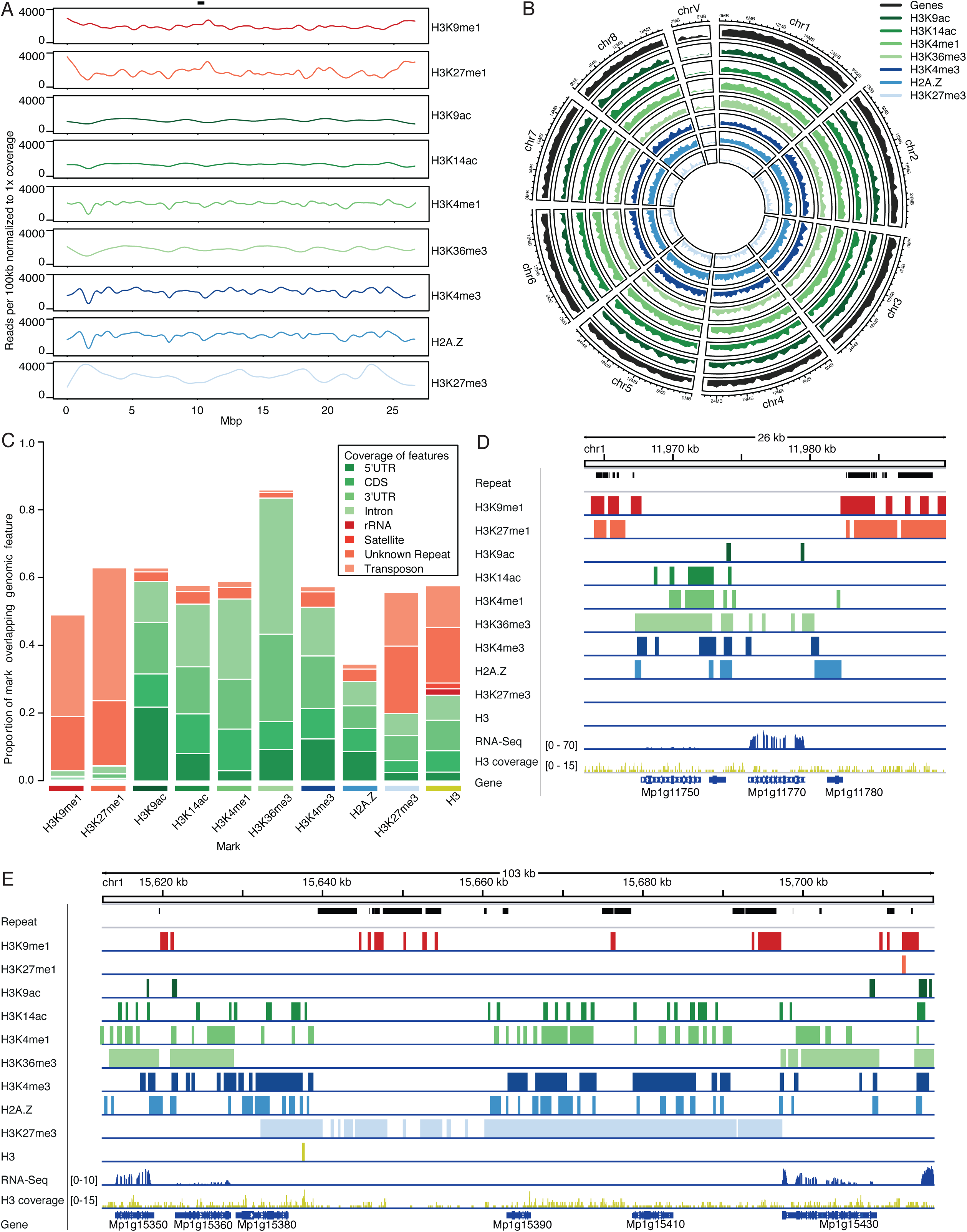
Distribution of chromatin marks in the *Marchantia* genome. (A) Coverage of chromatin marks across chromosome 5. Reads were normalized to 1x coverage and binned into 100kbp windows along the chromosome and a smoothed spline was fit to the data. Position of the putative centromere is indicated at the top. (B) Circos plot of euchromatic marks and genes. Each band shows the density of annotated chromatin mark peaks per chromosome, relative to the greatest density per band. (C) Distribution of chromatin marks over genomic features. The total length of chromatin mark peaks overlapping specified genomic features was divided by the total length of peaks of chromatin marks to determine each proportion. Unknown represents repeats annotated as unknown by RepeatMasker. Simple repeats not shown as they cover less than 0.3% of chromatin mark peaks. (D) IGV browser screenshot demonstrating flanking of genes by H3K9me1 and H3K27me1 marked transposons. The region shown is 26kb in length and from the proximal arm of chromosome 1. Chromatin mark tracks are bigwig files of peaks, except for the H3 coverage track which is a bigwig of mapped H3 reads. “Repeat” and “Gene” tracks are annotation files for repeats and genes, respectively. “RNA-seq” track is a bigwig of mapped RNA-seq reads from thallus tissue (Higo et al. 2016). (E) IGV browser screenshot demonstrating large H3K27me3 islands covering both genes and transposons. The region shown is 106kb in length and from the distal arm of chromosome 1. Tracks are as noted in (D).

### Histone modifications and gene expression

We explored preferential associations between chromatin marks and the transcriptional status of genes based on their average expression in the thallus somatic cells [10]. H3K36me3 showed the strongest association with expressed genes, which were also marked by H3K9ac, H3K14ac, and to a lesser extent by H3K4me1 and H3K4me3 (Figures 3A and S6A). In contrast, H3K9me1, H3K27me1, and H3K27me3 marked inactive genes (Figure 3A). Interestingly, H2A.Z showed a bimodal distribution of expression levels for the genes it associates with (Figure 3A), potentially linked with its correlation and overlap with H3K27me3 (Figure S5C).

**Figure 3.**
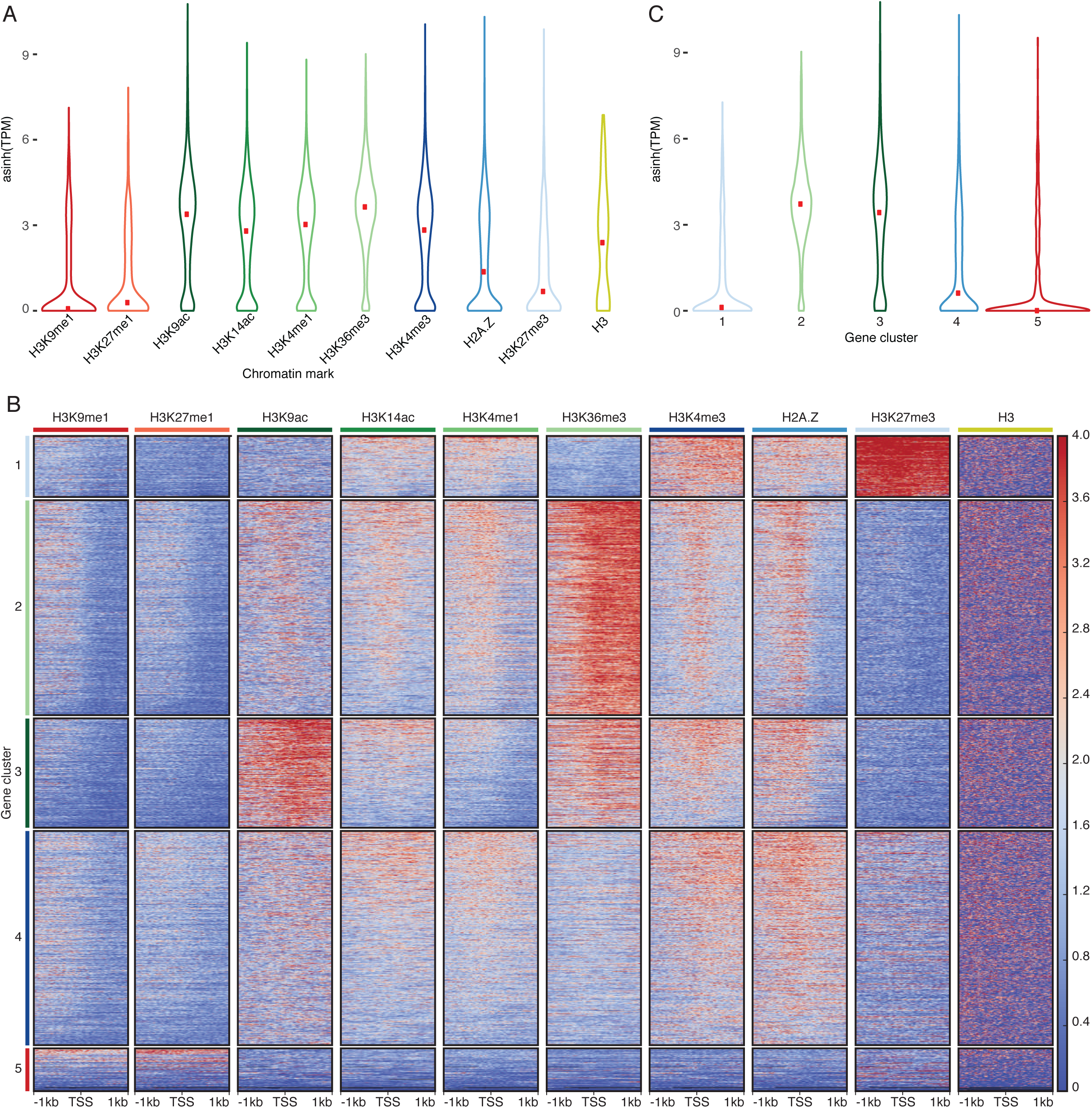
Association of chromatin marks with genes. (A) Expression level of genes associated with profiled chromatin marks. Width relative to density of genes. Red dots indicate median expression values. (B) Heatmap of k-means clustering of genes based on chromatin marks. Prevalence of each mark (columns) based on its z-score ±1kb around the transcription start site per gene, with red for enrichment and blue for depletion. Each row corresponds to one gene, with multiple genes grouped into blocks that have been defined as gene clusters 1 through 5. (C) Expression level of genes per gene cluster. Width relative to density of genes. Red dots indicate median expression values.

To untangle the relationships between chromatin profiles and genes in *Marchantia*, we performed k-means clustering of chromatin profiles over genes. This led to the identification of five main clusters of genes showing distinct chromatin environments (Figure 3B). Cluster 5 contained 7% of all genes and showed low levels of H3 and H3 modifications, suggesting a low nucleosome density, an inaccessibility for chromatin profiling, or difficulties in read alignment and we did not consider this cluster further. Gene clusters 2 and 3 encompassed active genes, accounting for 33% and 17% of genes, respectively, and showed enrichment in H3K14ac, H3K4me1, and H2A.Z at the TSS, though this trend was less marked for cluster 3 (Figures 3B, S6A and S6B). Genes in cluster 2 and 3 shared a strong enrichment in H3K36me3 over gene bodies with additional enrichment in H3K9ac in genes of cluster 3 (Figures 3B, S6A and S6B). Inactive genes were found in clusters 1 and 4, accounted for 10% and 33% of genes, respectively, and were characterized by a prominent enrichment of H2A.Z and H3K4me3 and an absence of H3K36me3 along gene bodies (Figures 3B, 3C, S6A and S6B). A strong enrichment of H3K27me3 distinguished genes in cluster 1 from genes in cluster 4 (Figures 3B and S6A). Gene clusters were uniformly distributed across the genome, to the exception of the gene-deprived sex chromosome V (Figure S6C). We observed a low density of DNA methylation in CG, CHG, and CHH contexts over genes irrespective of the nature of the dominating histone modification present (Figures S6D – S6F).

We conclude that DNA methylation on gene bodies does not correlate with chromatin states and transcriptional activity in *Marchantia* in contrast to *Arabidopsis* [34] and in agreement with a previous report [32]. In *Marchantia*, the enrichment in H3K36me3 over gene bodies is the best predictor of active transcription, and the combination of histone modifications that mark active genes is comparable to chromatin state 3 in *Arabidopsis* [4]. The TSS of active genes in *Marchantia* is marked by H3K4me3 and H2A.Z, similar to chromatin state 1, which marks TSS of active genes in *Arabidopsis* [4]. Repressed genes in *Marchantia* are marked with H2A.Z associated with H3K27me3 or H3K4me3 over gene bodies, similar to chromatin state 5 in *Arabidopsis* [4]. Altogether we conclude that how combination of histone modifications associate with gene transcriptional states in *Marchantia* is comparable to *Arabidopsis* [34], and other eukaryotes [35], although the association between H3K4me3 alongside H2A.Z on the body of inactive genes in cluster 4 appears more specific to *Marchantia*.

### Heterochromatin and transposons

We reassessed the census of transposons and repeats in *Marchantia*, which comprise at least 63 Mb representing 27% of the genome contrasting with 56% of the genome of *Physcomitrella* (Table S2). This lower proportion is largely attributed to the absence of the large expansion of Gypsy retrotransposons in *Physcomitrella* (Table S2 and [11]). In *Marchantia*, about two thirds of the transposons that were ascribed to a family belonged to retrotransposons from the Copia or Gypsy families and families of retrotransposons unique to *Marchantia* or *Physcomitrella* were identified (Figure 4A and Table S2). We also noted a comparable diversity of DNA transposons between the two species but an increased diversity of LINE families in *Marchantia* (Table S2), in part related to the expansion of LINE/RTE-X around centromeres (Figure S3C).

**Figure 4.**
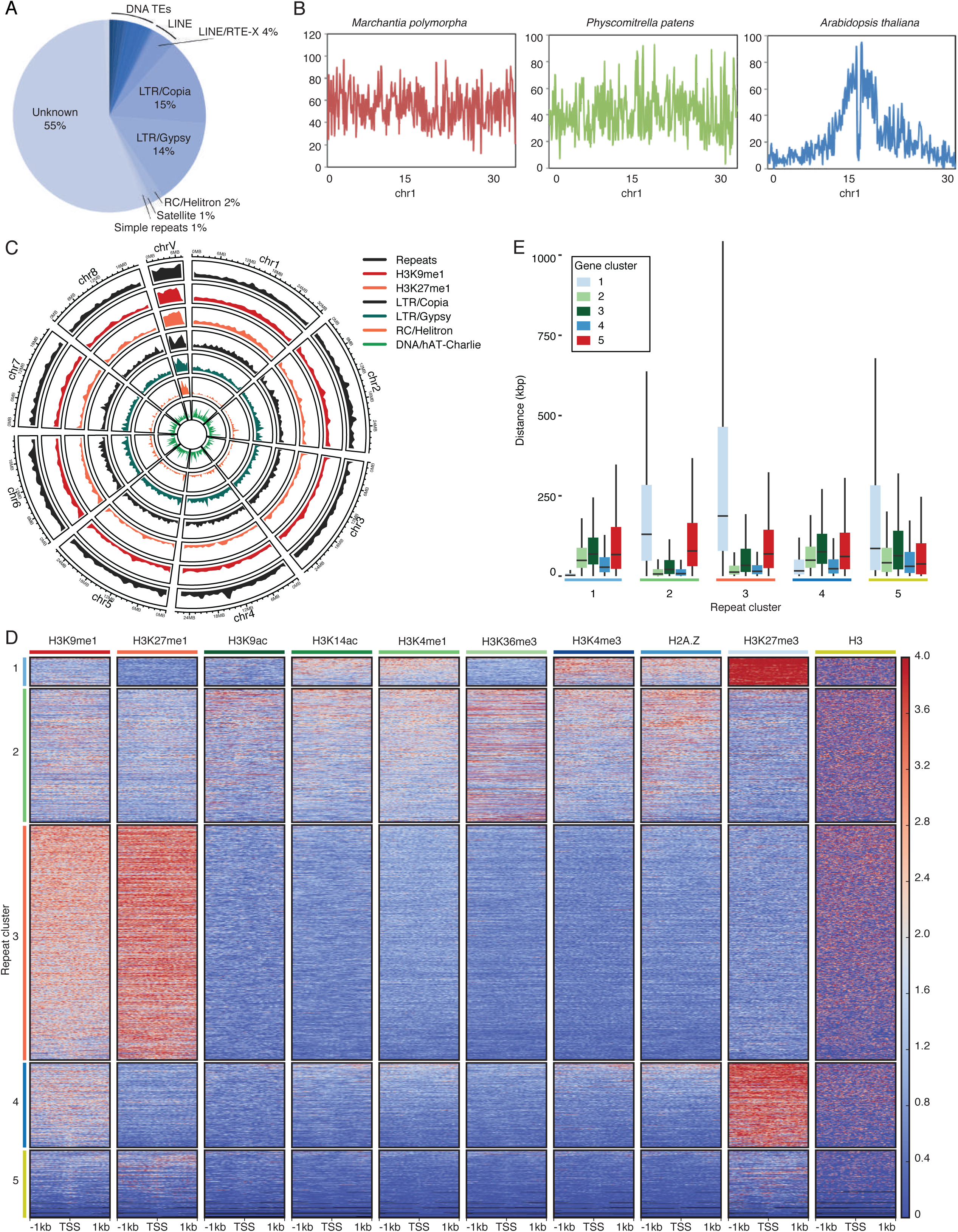
Association of chromatin marks with transposons. (A) Circos plot of heterochromatic marks, the four most abundant transposon superfamilies in *Marchantia* and all repeats. Each band shows the density of annotated repetitive elements or chromatin mark peaks per chromosome, relative to the greatest density per band. (B) Heatmap of k-means clustering of transposons based on chromatin marks. Prevalence of each mark (columns) based on its z-score ±1kb around the annotated start site per transposon, with red for enrichment and blue for depletion. Each row corresponds to one transposon, with multiple transposons grouped into blocks that have been defined as repeat clusters 1 through 5. (C) Boxplot of distances between each transposon and the nearest gene per gene cluster. Briefly each transposon is compared to all genes belonging to a gene cluster to find its nearest neighbor. Transposons are divided based on the repeat cluster they belong to. Distances in kilobases (kbp). Coloured boxes represent interquartile range and lines represent median values. Outliers not shown.

Heterochromatic marks and transposons were distributed evenly across chromosomes (Figures 4B and 4C). We performed k-means clustering of chromatin profiles over transposons and repeats leading to the identification of five main clusters showing distinct chromatin environments (Figure 4D). Over 40% of LINE/RTE-X elements were found in cluster 5 which represented 12% of repeats and was enriched around putative centromeres (Figure S3C). These transposons appeared to be relatively depleted of all profiled chromatin marks (Figure 4D), which could reflect a low nucleosome density or their relative inaccessibility to the MNase used in CUT&RUN profiling. Cluster 3, containing 43% of repeats and transposons, was characterized by a strong enrichment of H3K9me1 and H3K27me1 (Figures 4D and S7A). This cluster also associated with high DNA methylation levels in CG, CHG, and CHH contexts (Figures S7B – S7D) and the combination of chromatin marks in transposons and repeats from cluster 3 was comparable to chromatin states 8 and 9 in *Arabidopsis* [4]. Repeats from cluster 3 were much more enriched in the male sex chromosome V than on autosomes (Figures 4D and S7A). 25% of repeats and transposons represented cluster 2 that was enriched in DNA transposons (Figure S7E) and showed low uniform enrichment in all marks except H3K27me3 (Figure 4D). A similar chromatin state was observed over genes from cluster 4 (Figure 3B) and these two clusters were closely associated next to each other (Figures 4E and S7F). This combination of chromatin marks associated with low expression (Figure 3C) was not reported in *Arabidopsis*. Contrasting with clusters 2 and 3, H3K27me3 was enriched over transposons forming clusters 1 and 4, which represented 5% and 15% of repeats, respectively (Figure 4D). Repeats from cluster 4 showed higher levels of H3K9me1 whereas repeats from cluster 1 were more enriched in H3K4me3 and H2A.Z. DNA methylation levels in CG, CHG, and CHH contexts were higher in repeats from cluster 4 than from cluster 1 (Figures S7B – S7D). RC/Helitron elements were mostly enriched in cluster 4 whereas no major TE superfamily was enriched in cluster 1 (Figure S7E). Hence, we conclude that the clusters of repeats are not primarily differentiated based on the identity of the transposons and repeats or their position with the exception of the sex chromosomes that contain mostly repeats and transposons from cluster 3. These regions contrast with autosomes, where a large fraction of potentially mobile retrotransposons is marked by the repressive mark H3K27me3 (Figure S7E).

Strikingly, genes from cluster 2, which are expressed at high levels, were usually surrounded by transposons and repeats strongly enriched in H3K9me1 and H3K27me1 (Figures 2D and 4E). In contrast, H3K27me3 covered inactive genes and surrounding repeats and transposons (Figures 2E, 4E and S7F), accounting for 60% of nucleosomes that carried this mark related to the transcriptionally repressed state (Figures 2C and 3C). These account for large domains containing repressed genes and transposons covered by a high density of H3K27me3 (see an example in Figure 2E) in accord with potential of H3K27me3 to spread [36]. We conclude that a large proportion of genes and surrounding transposons share the same chromatin state in *Marchantia* with the notable exception being active genes surrounded by transposons marked by H3K9me1 on autosomes and exclusively so on the sex chromosome V.

### V chromosome and autosomes have distinct conformations

By comparing power-law decay curves of intra-chromosomal interaction strength with genomic distance in individual chromosomes, we found that the pattern of the male V chromosome was different from those of autosomes (Figures 5A and 5B). Particularly, the V chromosome Hi-C map indicated that it had stronger long-range chromatin contacts than those of autosomes, suggesting that the V chromosome was more compact. Additionally, on a chromosomal scale, the V chromosome exhibited significantly higher levels of heterochromatic marks H3K9me1 and H3K27me1 than autosomes (Figure 4C). These data indicate that the V chromosome is largely repressed and is more condensed than autosomes. Interestingly, manual inspection along the diagonal of the V chromosome Hi-C map revealed many self-interacting domains, in which chromatin contacts within one domain were stronger than those across different domains (Figure 5C). These self-interacting chromatin domains resembled topologically associated domains (TADs) discovered in mammals [37]. TADs appear as the basic structural units beyond nucleosomes, modulating higher-order chromatin organization [38]. TAD boundaries, which reflect local chromatin insulation, are enriched for insulator element binding proteins and active gene transcription [39]. Upon associating transcriptional activities at the V chromosome with the Hi-C map, we found a positive correlation in which many domain boundaries overlapped with local gene expression (Figure 5C). This suggests a tight relationship between the male sex chromosome topology and its transcriptional regulation.

**Figure 5.**
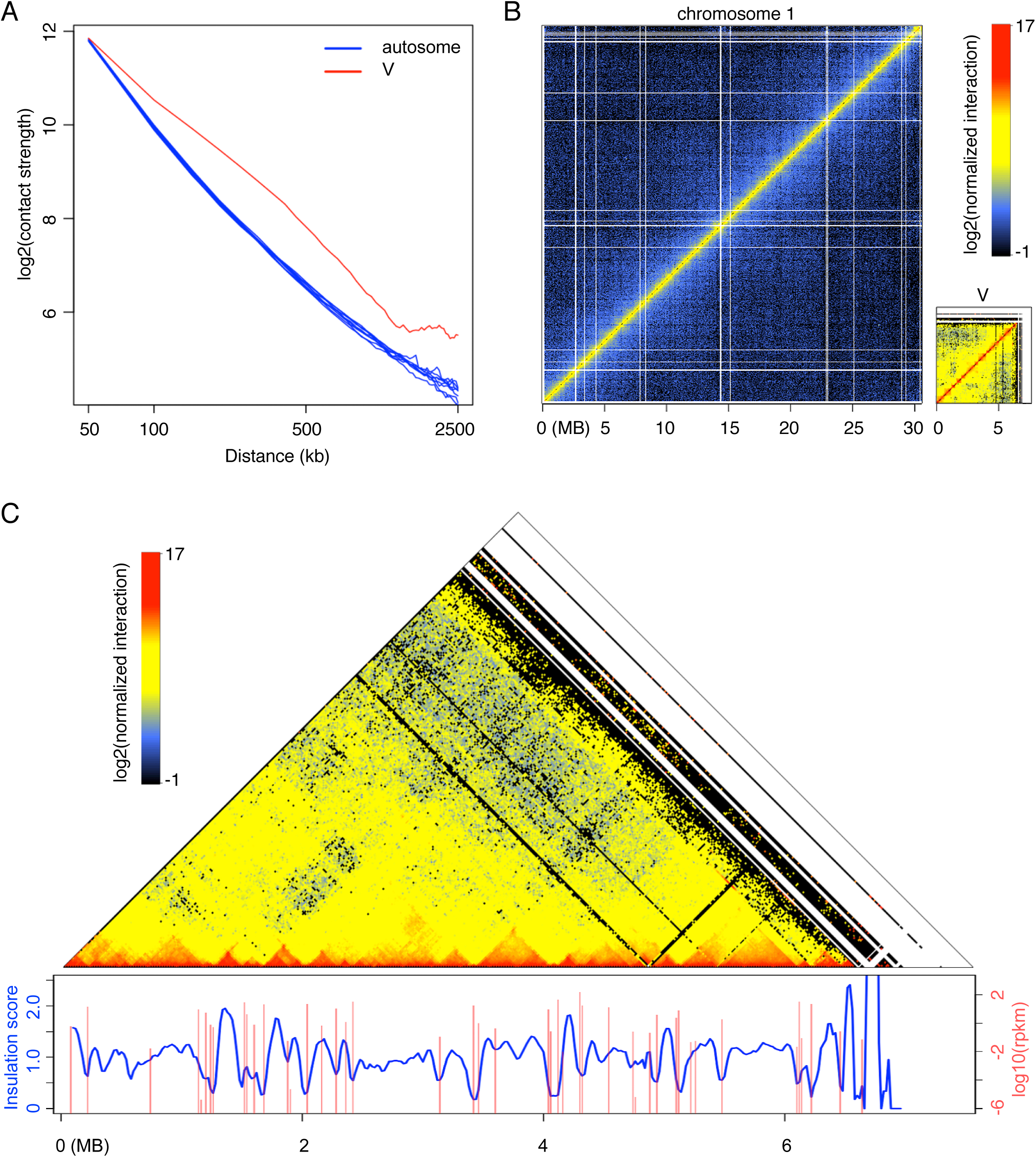
*Marchantia* chromosome V has distinct chromatin packing patterns compared with autosomes. (A) Comparison of interaction decay exponents among autosomes and V chromosome. The average interaction strengths of each chromosome at various distances were calculated based on a whole genome Hi-C map normalized at 50 kb resolution. (B) Hi-C maps of Tak-1 chromosome 1 and chromosome V. (C) Association between V chromosome Hi-C map (normalized at 20 kb resolution) and local gene expression. Insulation scores were calculated according to [65] with minor modifications, in which a sliding square of 100 kb x 100 kb along the matrix diagonal was used, and the ratio of observed over expected interaction strengths of this sliding square was plotted as insulation score. Genomic regions with local minima of insulation scores have strong chromatin insulation. Data of gene expression in Tak-1 thalli was from [10].

### Extensive intra- and inter-chromosomal contacts of *Marchantia* chromatin

On the genome-wide Hi-C map, we found many regions showing both strong intra- and inter-chromosomal contacts (Figure 6A). A comparison between interaction matrices generated with similar amounts of mapped reads from our Hi-C and a genome shotgun library indicated that these strong long-range chromatin interaction patterns were not caused by mapping errors (Figure 6B). Depending on their interaction networks, we classified these genomic regions into two groups (Figure 6C). One group (cluster 2) comprised regions found at chromosomal ends, consistent with our FISH data showing telomere clustering. This appears to be a universal phenomenon across plants [40–44].

**Figure 6.**
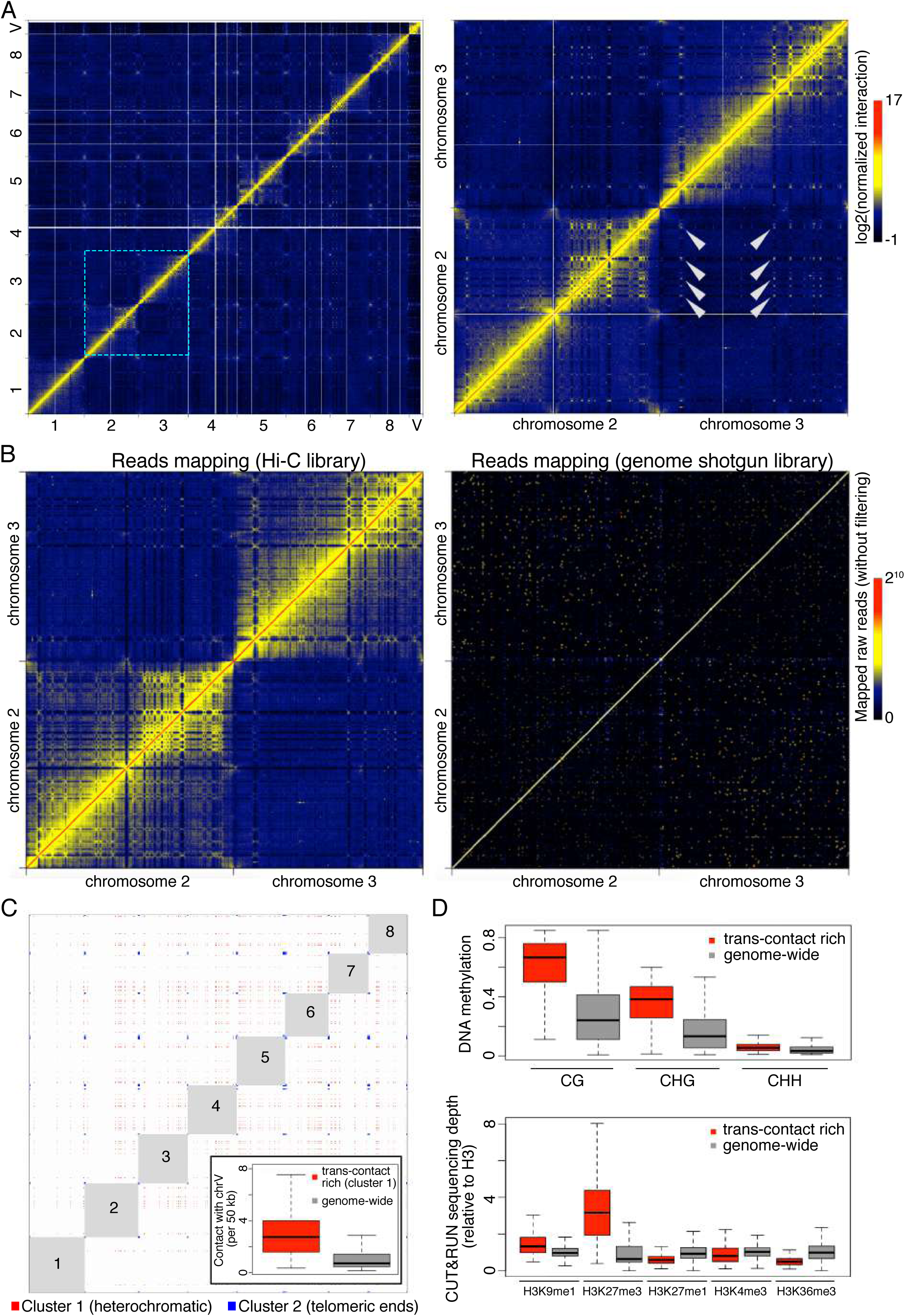
*Marchantia* genome shows extensive inter-chromosomal interactions. (A) Normalized Hi-C map at 50 kb resolution. The right panel shows the zoom-in image of an area containing chromosomes 2 and 3, in which selected trans-contacts among interstitial regions in different chromosomes are highlighted with arrowheads. (B) Comparison of chromatin interaction maps (50 kb bin) generated with comparable amounts of mapped reads in Hi-C and genome shotgun libraries (110 vs. 130 millions), respectively. The pair-end genome shotgun library is a combination of SRR396657 and SRR396658 [10], and was mapped to the assembled TAK-1 genome as Hi-C reads. Note that the diagonal of the plot shown on right has values larger than the maximum defined in the color bar. (C) Genomic regions showing strong and extensive trans-interactions. Bins having at least one top 0.5% inter-chromosomal contacts in the normalized Hi-C map shown in panel (A) were subjected to k-means clustering based on their genome-wide inter-chromosomal contact patterns. The optimal number of clusters was determined as 3 based on the Elbow method. For the first two clusters, virtual interactions among members of each cluster are shown as red and blue dots, respectively, representing an ideal situation in which all possible contacts happen within each cluster and are visible on a Hi-C map. Numbers depict autosome names. The inset shows inter-chromosomal contacts between autosomes and the V chromosome. (D) DNA methylation (top panel) and histone modifications (bottom panel) in genomic regions annotated as “cluster 1” in (C) and the whole genome (V chromosome not included). The DNA methylation data collected from Tak-1 thalli was from [32].

On the other hand, regions in the other group (cluster 1) were interstitial in each chromosome. Members of this group showed extensive contacts with each other, which stood out as speckles on the Hi-C map (Figures 6A and 6C, Table S3). These regions were depleted from the heterochromatic mark H3K27me1 and euchromatic marks H3K4me3 and H3K36me3 and showed enrichment in DNA methylation (Figure 6D). To some extent, these results resembled those associated with a special type of region in *Arabidopsis* and rice genomes named IHIs/KEEs (Interactive Heterochromatic Islands or KNOT ENGAGED ELEMENTs), which are marked by H3K9 methylation and DNA methylation [45–47]. In contrast with angiosperms, high levels of H3K27me3 were the strongest marker of heterochromatic islands in *Marchantia*. Notably, these heterochromatic islands showed stronger interactions with the V chromosome than did the average across all autosomes (Figure 6C, inset), suggesting the existence of chromatin compartmentalization that selectively brought some repressed genomic regions into physical proximity. Furthermore, a routine compartmentalization annotation to identify A (active) versus B (inactive) compartments [5] showed that B compartment regions were associated with trans-contact rich regions (Figure 7A). In contrast with A compartments marked by a strong association with H3K36me3, B compartments showed the highest levels of H3K27me3 and no significant association with enrichment in H3K9me1 and H3K27me1 (Figure 7B). We speculate that H3K27me3 plays an important role in shaping chromatin compartmentalization and defining heterochromatin in autosomes while local transcriptional activities delimit TADs on the sex chromosome.

**Figure 7.**
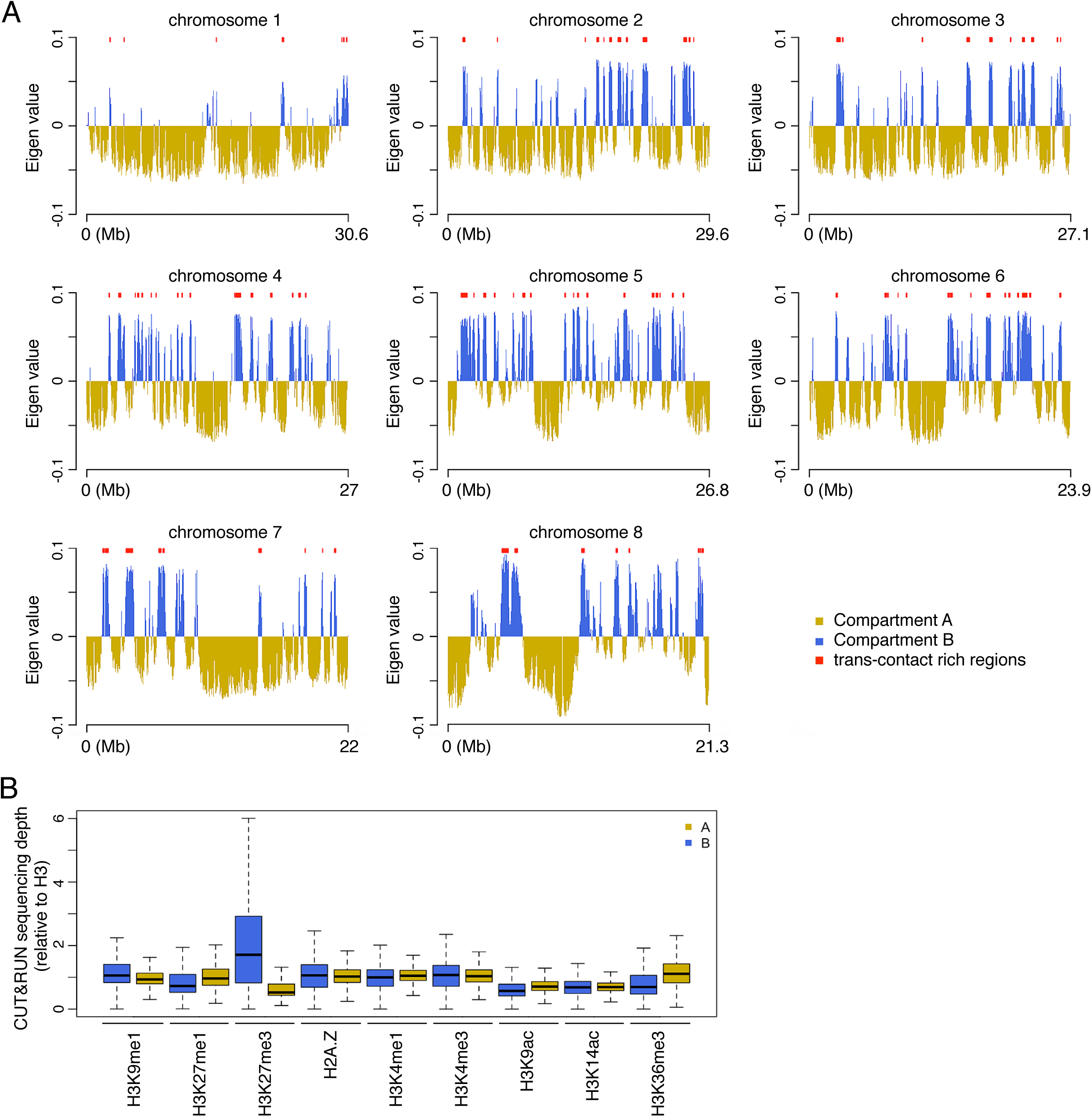
A/B compartments and their associated epigenetic marks. (A) A/B compartments in individual Tak-1 autosomes. For each autosome, the compartment bearing the estimated centromere is labeled as “Compartment B”. Red segments above each plot denote trans-contact rich region that display strong inter-chromosomal interactions. (B) Epigenetic features associated with A/B compartments.

## DISCUSSION

In flowering plants, transposons represent 10 to 90% of genomes and tend to cluster in pericentromeric heterochromatin clearly delimiting chromocenters, as shown in *Arabidopsis* [22, 24, 25]. In contrast, transposons and genes are spread relatively evenly across chromosomes in the moss *Physcomitrella patens* [11] and the liverwort *Marchantia polymorpha*. This is associated with the lack of chromocenters in both species and many other bryophytes including hornworts [48], suggesting that early land plants shared a general genome organization devoid of a linear cluster of transposons. It has been proposed that the interspersed organization of genes and transposons in *Physcomitrella* may be a facet of inbreeding and low recombination rates [11]. As *Marchantia* and many other liverworts are dioicous and reproduce by outcrossing, there are likely alternative explanations. However, the enrichment of specific classes of transposons around the centromeres of *Physcomitrella* and *Marchantia* indicates that potential mechanisms by which transposons become enriched around centromeres may have been active already in these plants.

Epigenetic and transcriptional states are key predictors of Hi-C contact maps in eukaryotes [39, 49, 50]. Similar to the observations made from Hi-C maps in other eukaryotes, the binary annotation of *Marchantia* autosomes based on Hi-C data largely correlates to the demarcation of active/inactive chromatin domains. On the V chromosome, DNA and H3K9 methylation are associated with transposons surrounding highly expressed genes, forming clear topologically associated domains. These associations also exist on autosomes (Figure 2D) but are relatively scarce compared with the sex chromosome. Similar patterns are also observed in *Arabidopsis* chromocenters, in which the 3D folding of constitutive heterochromatin marked by DNA and H3K9 methylation is proposed to be driven by local expression levels [39]. This suggests that the function of marks typical of constitutive heterochromatin in eukaryotes [51] is conserved in *Marchantia* and insulates transcriptional units.

The majority of the *Marchantia* genome exhibits low levels of DNA methylation [32], as in other bryophytes [52, 53], and we observed that a large fraction of transposons and repeats are not marked by H3K9me1 and H3K27me1. In *Marchantia*, these marks do not associate with repressive B compartments and trans-contact rich regions, whereas these type of regions represent constitutive heterochromatin marked by H3K9me1 and H3K27me1 in flowering plants [54]. Remarkably, half of transposons are marked with H3K27me3. H3K27me3 is deposited by the Polycomb repressive complex 2 (PRC2) in *Physcomitrella* [55] and the conservation of PRC2 subunits in *Marchantia* [10] indicates that its function is likely conserved in bryophytes. In land plants, as in other eukaryotes, H3K27me3 is involved in maintaining repressed transcriptional states [4, 55, 56] and previous plant Hi-C studies reported that H3K27me3-marked chromatin is involved in forming long-range interactions [46, 57, 58]. Hi-C analyses in *Marchantia* highlight the dominant impact of H3K27me3 in strong intra- and inter-chromosomal contacts. The heterochromatic islands marked by H3K27me3 in *Marchantia* are likely to be distinct from heterochromatic islands marked by H3K9 methylation in flowering plants both in their genesis and association with transcriptional regulation. H3K27me3 forms domains along the linear genome comprising genes and transposons. This contrasts with flowering plant transposons that associate primarily with H3K9me2 [4], although in *Arabidopsis*, a fraction of transposons are marked by H3K27me3 in reproductive tissues which are characterized by reduced DNA methylation [59] and in mutants with reduced DNA methylation (Bioarchive **doi:** https://doi.org/10.1101/782219). We thus propose that PRC2 targeted deposition of the repressive mark H3K27me3 on transposons in the ancestors of land plants. In *Marchantia*, the association between H3K27me3 and transposons is still extant. This might be explained by the absence of a strong feedback loop between DNA and H3K9 methylation in bryophytes [60]. The association between a few transposons and H3K27me3 has been reported in red algae, a group that diverged from the streptophyte lineage more than 900 Mya [61] and phylogenetic data support the emergence of PRC2 function in unicellular eukaryotes [62]. In ciliates H3K27me3 is also associated with transposons silencing, where it is deposited together with H3K9me3 by PRC2 [63]. In contrast, we observe a clear distinction between the group of transposons marked by H3K9 methylation and H3K27me3 in *Marchantia*, which may result from the PRC2-independent evolution of the H3K9 methylation pathway in plants [2, 60, 64]. It remains to be investigated whether H3K27me3 led to transposon silencing in ancestors of land plants and *Marchantia* appears to be an ideal model for such studies.

## Acknowledgements

We acknowledge computing support by the High Performance and Cloud Computing Group at the Zentrum für Datenverarbeitung of the University of Tübingen, the state of Baden-Württemberg through bwHPC and the German Research Foundation (DFG) through grant no. INST 37/935-1 FUGG. We acknowledge Ms. Fumi Hayashi and Dr. Mika Sakamoto for helping exhaustive manual correction of the assembly. FB acknowledges support from the PlantS, next generation sequencing and histopathology facilities at the Vienna BioCenter Core Facilities (VBCF), and the BioOptics facility and Molecular Biology Services from the Institute for Molecular Pathology (IMP), and Dr. J. Matthew Watson for proof-reading the manuscript.

CL and NW were supported by European Research Council (ERC) under the European Union’s Horizon 2020 research and innovation programme (grant agreement No. 757600). This work was also supported by the Gregor Mendel Institute (FB and SA) and FWF grants I2163-B16, I2303-B25, P26887, and DK 1238 chromosome dynamics (SAM and FB); National Institutes of Health (R01 GM065383 to DES.; R01 GM127402 to EVS), Russian Science Foundation (18-74-00112 to LRV), Russian Foundation for Basic Research (18-016-00146 to EVS) and funds from the Russian Government Program for Competitive Growth of Kazan Federal University. JSPS KAKENHI grant numbers 16H06279 (YT, YN, and TK), 15K21758 (TK, FB, and YN), 17H05841 (SY), 25113001 (TK) and 25113009 (TK); the Project Research of the Faculty of Biology-Oriented Science and Technology, Kindai University No. 16-I-3,2017 (KTY), the Australian Research Council, DP170100049 (JLB).

## Authors contributions

SA produced the DNA for PacBio sequencing, CL, BG, and YT performed the genome reassembly with help provided by YN and TY. SAM analyzed chromatin modifications and epigenetic landscapes. CL performed all Hi-C analyses. YT, TM, and MY performed the gene annotation, TI performed the TE annotation. Telomere analysis was performed by LRV, EVS, and DES. Centromeres were defined by SAM and further identified in microscopy by NW, who also performed the cytogenetic analysis of telomeres. tRNAs were analyzed by VC and LD, miRNAs were analyzed by SSL. TK laboratory contributed CAGE data that was analyzed by SY. Iso-seq data was obtained by KTY. FB and CL conceived the project. FB, CL, and SAM wrote the manuscript. YT and YN conceived the webpage interface and handled data repository at NIG.

## METHODS

### MATERIALS AND METHODS

#### Plant Material

Male Takaragaike-1 (Tak-1) [66] (*Marchantia polymorpha*) gemmae were cultured on half-strength B5 medium supplemented with 1% sucrose. The light condition was set to long day (16 hr light and 8 hr dark, 3,000 lux) and the temperature was maintained at 22 °C.

#### Isolation of nuclear DNA from *Marchantia*

Briefly, 100 g of 3-week-old thallus was rinsed with 250 mL of ice-cold ethyl ether for 3 minutes followed by washing with cold TE buffer, and homogenized with 1 L of cold MPD-based extraction buffer (1 M 2-methy-2,4-pentanediol, 10 mM PIPES-KOH, 10 mM MgCl2×6H2O, 2% polyvinylpyrrolidone (PVP), 10 mM sodium metabisulfite, 5 mM 2-mercaptoethanol, 0.5% sodium diethyldithiocarbamate, 200 mM L-lysine, and 6 mM EGTA, pH 6.0.). The slurry was filtered through a 40 µm nylon filter, and Triton X-100 was added to the flow-through to 0.5% v/v. The mixture was centrifuged at 800 x g for 20 min at 4°C, and the nuclei pellet was washed three times with MPDB buffer (0.5 M 2-methy-2,4-pentanediol, 10 mM PIPES-KOH, 10 mM MgCl2×6H2O, 0.5% Triton X-100, 10 mM sodium metabisulfite, 5 mM 2-mercaptoethanol, 200 mM L-lysine, and 6 mM EGTA, pH 6.0.). Nuclei were then lysed with 2% SDS (w/v) at 60°C for 10 min, and the released genomic DNA was extracted with phenol/chloroform/isoamyl alcohol (25:24:1) following the standard protocol. The aqueous layer was dialyzed overnight into TE buffer at 4°C. On the next day, RNase T1 and RNase A were added to the sample to a final concentration of 50 units/ml and 50 µg/ml, respectively. RNA digestion was performed at 37°C for 60 min. Subsequently, Proteinase K was added to a final concentration of 150 µg/ml, and the solution was further incubated at 37°C for 60 min. Finally, DNA was recovered by following standard phenol/chloroform/isoamyl alcohol extraction and ethanol precipitation protocols.

#### Hi-C library preparation and sequencing

The *in situ* Hi-C library preparation was performed by following a protocol established for rice seedlings [41] In total, two replicates of 3-week old Tak-1 thalli Hi-C libraries were made, and for each replicate around 0.5 g of fixed sample was homogenized for nuclei isolation. The libraries were sequenced on an Illumina HiSeq 3000 instrument with 2 x 150 bp reads.

#### Chromosome-scale genome assembly

PacBio reads were assembled into scaffolds with miniasm using default settings [67] except that the minimum coverage was set as -c 2. Next, Hi-C reads were mapped to these scaffolds with an iterative mapping strategy described previously [41]. Subsequently, Hi-C contacts were processed by the 3d-dna-master software to further assemble the scaffolds [68]. In brief, the whole process had two steps. Firstly, it attempted to connect all scaffolds to build a genomic “super-scaffold”. Next, it split this “super-scaffold” into chromosomes according to the chromosome number defined by the user. For the first step, a Tak-1 “super-scaffold” was generated with following parameters: -t 1000 -s 3 -c 9 -w 25000 -n 1000 -k 5 -d 150000. Consistent with Tak-1’s karyotype, this “super-scaffold” showed 9 blocks of self-interacting domains with various sizes (Figure S1) [69]. For the second step, we split this “super-scaffold” into 9 segments (chromosomes) with the parameter set as -c 9 accordingly. Because the estimated size of the Tak-1 V chromosome (10 Mb) is much smaller than the minimum expected chromosome size to be split from the “super-scaffold” by the 3d-dna-master program, we modified two default settings to circumvent this issue [15]. We changed the resolution setting (“res”) in the “run-asm-splitter.sh” file from 100000 (default) to 50000, and the bin number setting (“m_size_threshold”) in the “recursive-chromosome-splitter.py” file from 200 (default) to 60. In this way, we modified the lower boundary of “chromosome size” that the program accepted to 3 MB (50000 kb x 60), which is smaller than that of the V chromosome. As a result, the 3d-dna-master tool generated an assembled Tak-1 reference with 9 “chromosomes” that collectively covered around 215 MB as well as 441 unplaced scaffolds adding up to 3 MB that failed to be localized to any chromosomal sequence.

Next, we manually searched for local misjoint errors by checking the diagonals of Hi-C maps at 20 kb window setting. Typically, mapping Hi-C reads to a reference containing misjoints or large-scale chromosomal rearrangements gives rise to aberrant and strong “interactions” off the diagonals in Hi-C maps. Meanwhile, these regions display depleted interactions with their neighboring chromatin (see examples in Figures S1B and S1C, left panels). Upon identifying misjoints, we rearranged the corresponding scaffolds according to the Hi-C map such that the revised scaffold ordering would generate a continuous diagonal (Figures S1B and S1C, right panels). Finally, the manually inspected and corrected chromosomes were sorted in descending order according to their size and named chromosome 1 to 8 and V.

#### Genome assembly polishing

The chromosome-level assembly of the Tak-1 genome was further processed with the Pilon tool for local sequence correction [70]. A subset of Illumina short reads from Tak-1 (SRR1800537), which correspond to approximately 100X genomic coverage, were preprocessed using fastp with “--cut_front --cut_tail” options. They were aligned to the pre-polished Hi-C assembly using BWA v0.7.15 with the MEM algorithm. The alignment result was provided to Pilon version 1.22 to correct short indels and SNPs (--fix indels,snps). Additionally, indels and SNPs in the protein-coding regions were corrected manually based on the mapping results of RNAseq and Iso-seq.

#### Gap closing and additional scaffolds

Assembly gaps in the polished genome sequences were filled with the version 3.1 (v3.1) sequences after checking the flanking regions and the order of protein-coding genes within and around the gap. When both of the flanking 800 bp regions of the gap matched with v3.1 sequences (>99% identity) and the gene order was consistent when compared to the annotation in the v3.1 genome, the gap was fully patched with the v3.1 sequence. When only one of the flanking 800 bp regions matched the v3.1 sequence, the gap was partially patched with the v3.1 sequence containing the target genes. In total, 52 assembly gaps were fully patched and 32 were partially patched.

When gene sequences from v3.1 genome, whose annotation was well supported by expression evidence and/or protein homology, were not mapped to the assembled genome, genomic regions containing those v3.1 genes were added as unplaced scaffolds. This resulted in additional 14 scaffolds. 20 unplaced scaffolds were removed from the assembly as they were redundant or considered to be derived from chloroplast genomes. We finally obtained the genome assembly designated as v5.1, which consists of 9 chromosomal sequences and an additional 435 unplaced scaffolds.

#### CAGE-seq, Iso-seq, and data analysis

CAGE-seq and Iso-seq were employed for improving gene annotation. For CAGE-seq analysis, total RNA was isolated with an RNeasy kit (QIAGEN) from 10 day-old Tak-1 thalli cultured from gemmae under continuous white fluorescent tube light. CAGE library construction, sequencing, and mapping onto the v5.1 genome was carried out by DNAFORM (Yokohama, Kanagawa, Japan). The mapped read distribution on the v5.1 genome was calculated by RSeQC ver.3.0.0 [71]. For Iso-seq analysis, total RNA was separately prepared by an RNeasy kit from the meristematic regions of 10 day-old thalli cultured from gemmae (vegetative tissue) and immature gametangiophores (reproductive tissue) for each of Tak-1 (male) and Tak-2 (female) plants, and then pooled to make male and female pooled samples, each of which contains RNA from two different tissues. Library construction and sequencing by PacBio Sequel (Pacific Biosciences, Menlo Park, CA, USA) were carried out by Kazusa DNA Research Institute (Kazusa, Chiba, Japan). Obtained data were processed with the IsoSeq3 pipeline of SMRT Link v6.0 (Pacific Biosciences) to generate clean sequences and they were aligned to the genome using GMAP (ver. 2018-07-04)[72].

#### Genome annotation

Annotation of protein-coding genes was conducted through a combination of the ver 3.1 genome and *de novo* prediction. A total of 24,674 predicted transcript models (including 5,387 isoforms) for the v3.1 genome were obtained from MarpolBase (http://marchantia.info). After excluding 134 genes putatively encoded on the female sex chromosome, they were aligned to the v5.1 genome sequences using BLASTN. The 23,623 transcript models (96.2%) that were aligned without insertions or deletions within coding regions were transferred from the v3.1 genome. Subsequently, 455 were aligned to the v5.1 genome with GMAP and manually modified if needed. The remaining 462 transcript models, which were not supported by expression data or protein homology, were discarded as false genes.

For *de novo* gene prediction, RNA-seq libraries (SRR896223-30, PRJNA251267) were mapped to the repeat-masked genome using Hi-SAT2 (ver. 2.1.0) [73]. The mapping results were used to build transcript models using Braker2 (ver. 2.0.3) [74] and StringTie (ver. 1.3.4d) [75]. Braker2 was run with the Augustus parameters pre-trained using ver. 3.1 gene models. In total, 166 and 89 transcript models were incorporated from the results of Braker2 and StringTie, respectively. Based on manual inspection using RNA-seq and Iso-seq, 418 transcript models were also added. Functional annotation for transcript modelling was performed by an RPS-BLAST search against the Eukaryotic Orthologous Groups (KOG) database [76], KEGG pathway analysis using KEGG Automatic Annotation Server (KAAS) [77], and InterProScan [78].

The completeness of the gene set was evaluated by BUSCO using 303 universal single-copy orthologous markers designed for eukaryotes (eukaryota_odb9) [14].

Repeat masking was conducted using RepeatModeler (ver 1.0.11) and RepeatMasker (ver. 4.0.7) (http://www.repeatmasker.org). A *de novo* repeat library was constructed using RepeatModeler, which was then subjected to RepeatMasker as a custom library to mask repetitive regions of the genome. RepeatMasker was run with ‘-s -no_low’ parameters.

The annotation of micro-RNA genes and their putative targets was based on published information [79, 80].The mature miRNA and v5.1 mRNA profiles were used for putative target prediction by psRNATarget [81]. The degradome profile from Tak-1 thallus (SRR2179617) was used to evaluate the target prediction based on the method that was published previously [79]. Putative targets had to fit the following criteria: (1) degradome reads of the cleaved site (CS-d reads) had to be greater than or equal to 5 reads; (2) the CS-d read count was claimed significant larger than the nearby 100 bp window (±50 bp from the site) if the p-value of Poisson one-tail test was less than 0.05. Details of miRNA sequences and their target gene identities can be found in Table S2.

Nuclear tRNA prediction was done with tRNAscan-SE version 2.0 using the general model parameter [82]. The data were manually curated to filter tRNA, organellar contaminations, and tRNA-like sequences. Details of each nuclear tRNA locus can be found in Table S2.

Large sequence comparison of sex chromosomes from v3.1 and v5.1 were aligned and visualized by D-Genies with default parameters [83].

#### Chromatin profiling and data analysis

*Marchantia* Tak-1 gemmae were cultured on half-strength B5 medium under continuous light at 22°C for 14 days. Plants, excluding gemmae cups, were chopped in Galbraith buffer (45 mM MgCl_2_-6H_2_O, 30 mM Trisodium citrate, 20 mM MOPS) pH 7.0 plus 0.1% Triton-X 100 with a razor blade on ice to extract nuclei. Nuclei were passed through a 40 μm filter and stained with 2 μg/mL DAPI before sorting on a BD FACSARIA III (BD Biosciences). Aliquots of 40,000 nuclei were collected in 10X binding buffer (200 mM HEPES-KOH pH 7.9) diluted 1:10 in 1x PBS. The harvested nuclei were processed with the CUT&RUN protocol [31].

CUT&RUN reads were mapped to the Tak-1 v5.1 genome presented in this paper using Bowtie2 v2.1.0 [84] and further processed using Samtools v1.3 [85] and Bedtools v2.17.0 [86]. Reads with MAPQ less than ten were removed with Samtools v1.3 and duplicates were removed with Picard v1.141 (http://broadinstitute.github.io/picard/). Inserts less than 150 bp were removed from further analyses, as these fragments are sub-nucleosomal in size and likely represent noise when profiling histones and histone modifications. Deduplicated reads from 2-4 biological replicates were merged. We called peaks for chromatin marks using HOMER [87] and considered a gene associated with a mark if at least 50% of the gene length overlapped with peaks. We used the following settings: -style histone -size 250 -minDist 500. Bigwig files were made using deepTools v2.2.4 [88].

Pearson correlation matrices were generated using deepTools v2.5.4 [88] using multiBamSummary and plotCorrelation tools. Overlaps between features were calculated using bedtools intersect v2.27.1 [86]. Circos plots were generated using circlize [89] using bedgraphs of peaks called by HOMER. Chromosome coverage plots were generated using the smooth.spline function in R v3.4.0 (https://www.R-project.org/). IGV v2.3.97 [90] browser shot was obtained by loading bed files of peaks and bigwig files of RNA-Seq and H3 coverage data.

#### Gene expression analyses

Gene expression data from [91] were downloaded from the SRA (samples DRR050343, DRR050344, DRR050345) and processed with RSEM v1.2.31 [92] and STAR v2.5.2a [93]. Transcript Per Million (TPM) values were averaged from three biological replicates from vegetative thalli and used for further analyses. Genes were determined to overlap with a feature of interest if at least 50% of the gene length overlapped with the feature.

#### Clustering analyses

K-means clustering of chromatin marks was performed using deepTools v2.2.4 [88]. Matrices were computed using computeMatrix for either genes or repeats using bigwig files as input and the start of the feature as the reference point with 1 kb upstream and downstream. Heatmaps of matrices were plotted with plotHeatmap with k-means clustering. Cluster assignments can be found in Table S5.

#### DNA methylation analysis

Bisulfite sequencing data of Tak1-1 thallus was downloaded from SRA (SRP101412) and analyzed following the method described in [32]. Read mapping and the identification of methylated cytosines were performed with Bismark v0.22.1 with default settings [94]. The mean methylation percentage per gene or repeat was calculated using MethylDackel v0.4.0 (https://github.com/dpryan79/MethylDackel) from analyzed cytosines that were assigned to genes or repeats.

#### Nuclei immunostaining

*Marchantia* Tak-1 thallus and *Physcomitrella patens* gametophyte were chopped in Galbraith buffer (45 mM MgCl_2_-6H_2_O, 30 mM Trisodium citrate, 20 mM MOPS) pH 7.0 plus 0.1% Triton-X 100 with a razor blade on ice to extract nuclei. Nuclei were passed through a 40μm filter and immunostained following a protocol by [95]. Images were obtained on an LSM 780 (Zeiss) and processed using FIJI [96]. Images shown are maximum intensity projections. Contrast was enhanced for *Marchantia* H3K27me1 and H3K27me3 stainings and *Physcomitrella* H3K4me3, H3K27me1, and H3K27me3 stainings.

#### Hi-C map normalization

Raw Hi-C reads of the two replicates used for genome assembly were mapped to the final Tak-1 genome assembly. Read mapping and filtering were performed essentially as described [41]; at the end, about 89 million informative Hi-C reads were obtained in total (Table S4). Hi-C matrices normalization was performed as described [41] assuming equal visibility of individual genomic bins, with which a Hi-C matrix was adjusted towards having similar sum values for each row or column [97]. Normalization of the Hi-C map at 50 kb resolution was performed at the genome-wide level (i.e., all chromosomes were included), while normalization at 20 kb was done separately for each chromosome.

#### Chromosome spread preparation and Fluorescence in situ Hybridization (FISH)

Chromosome spread preparation was performed as described [16] and placed on Superfrost Ultra Plus Adhesion Slides (ThermoFisher Scientific). Centromeric repeats probes were synthesized as two oligos: 5’-[DIG]TGGGCTTGTTCACGACGGCCGGGCGCACATACCTGCAAATTTTCAGCCCCAACGGAGCT[DIG]-3’ and 5’-[DIG]TTTTCAGCCCCAACGGAGCTGCTGTCAAGAAGTTGTCATTTCGAAACTTTGAGTTT[DIG]-3’, (Figure S3B) where the terminal thymidines were labeled with digoxigenin (DIG). These two oligos were mixed in a 1:1 molar ratio and used for hybridization. Telomere probes were synthesized as 5’-[DIG](TTTAGGG)_7_T[DIG]-3’. For probe hybridization, 5 µl of hybridization buffer [54] containing 25 ng DIG-labeled telomere probes was used. Before applying the probes to the slides, the probes were denatured at 95°C for 5 min and cooled for 5 min on ice. For hybridization, the slides were heated at 70°C for 8 min and incubated at 37°C overnight in a humid chamber. Detection of the DIG probes was performed according to [54].

For FISH experiment with *Marchantia* nuclei, around 5,000 nuclei were collected with FACS as described [98] and were used for one hybridization spot (∼ 1 cm^2^). After nuclei sorting, the nuclei were centrifuged for 3,000 x g at 4°C for 7 min, and the pellet was resuspended with 20 µl PBS buffer. The nuclei were incubated at 65°C for 30 min, and mixed with 5 µl 0.1 mg/ml RNase A. The mixture was transferred onto a Superfrost Ultra Plus Adhesion Slide (ThermoFisher Scientific) and incubated for 1 h at 37°C. At the end of RNase A treatment, the nuclei became attached to the glass slide. Next, the slide was washed briefly with PBS buffer and dehydrated in a graded series of alcohol solutions. All subsequent steps, including probe denaturation, hybridization, washing, and detection were performed as described for chromosome spread samples.

#### Centromere identification

Regions with strong Hi-C interactions amongst each other and occurring only once per chromosome were aligned to create dot plots using EMBOSS Dotmatcher with 10 bp windows and a threshold of 50 [99] (Figure S3D). One 165 bp repeat found in each region was identified and the centromeric FISH probes are indicated (Figure S3B).

#### Data availability

All raw read data and assembled sequence data that support the findings of this study have been submitted to the DDBJ/ENA/NCBI public sequence databases under accession numbers PRJNA553138 and PRJDB8530.

**Supplemental Figure 1.**
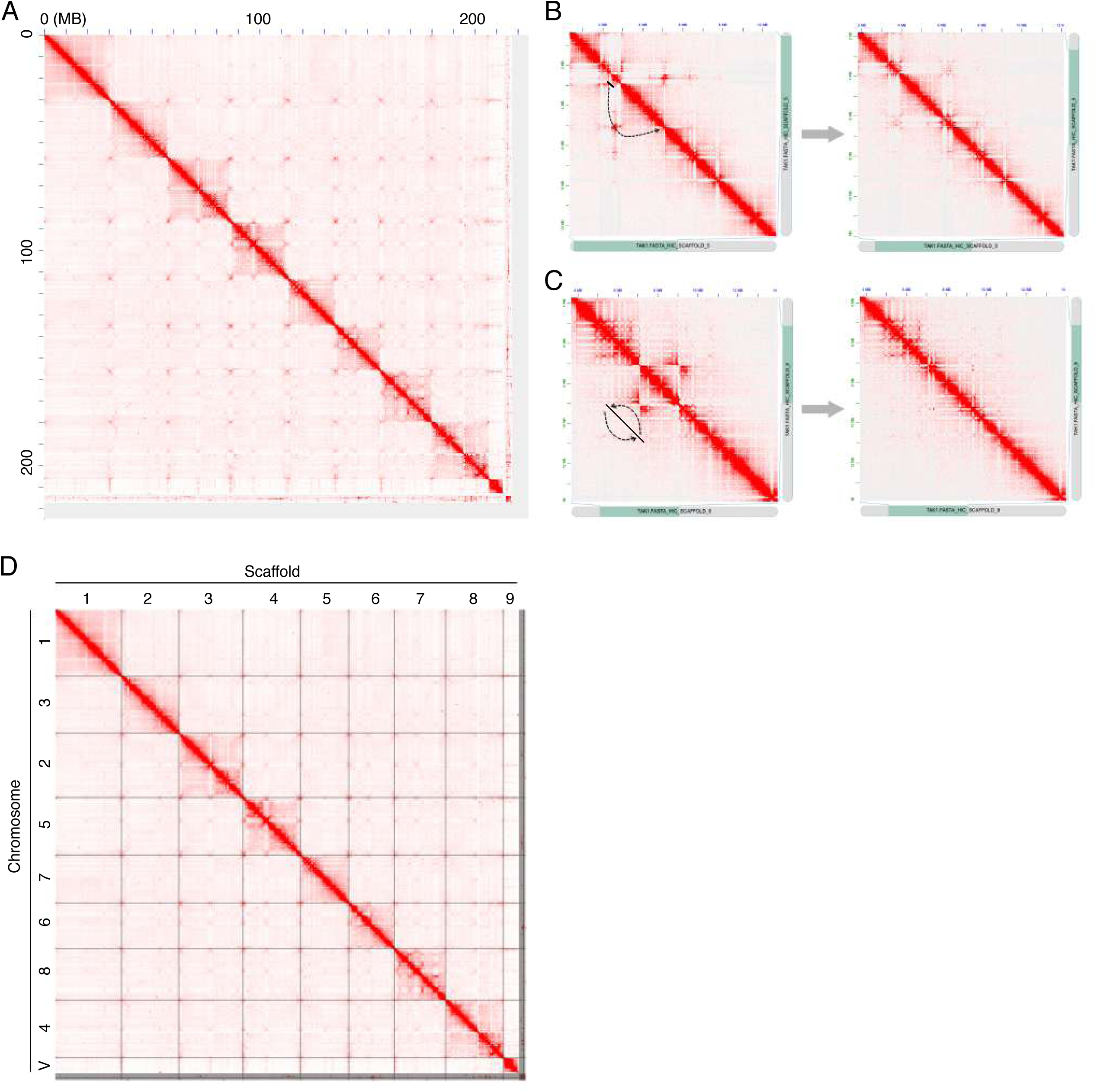
Hi-C guided assembly of the *Marchantia* Tak-1 genome. (A) Hi-C map of the assembled “super-scaffold”, visualized with Juicebox [1]. The vast majority of this super-scaffold is consisting of 9 distinct self-interacting blocks. (B and C) Manual inspection and correction of local misjoints. Panels on right show corrected Hi-C maps. Depending on the nature of aberrant interaction patterns, they can be corrected by changing the order of scaffolds, such as shift (B), inversion (C), or a combination of them. (D) Hi-C map of the Tak-1 genome with manual correction. The nomenclature of chromosomes 1 to 8 and chromosome V is according to the sizes of the longest assembled 9 scaffolds.

**Supplemental Figure 2.**
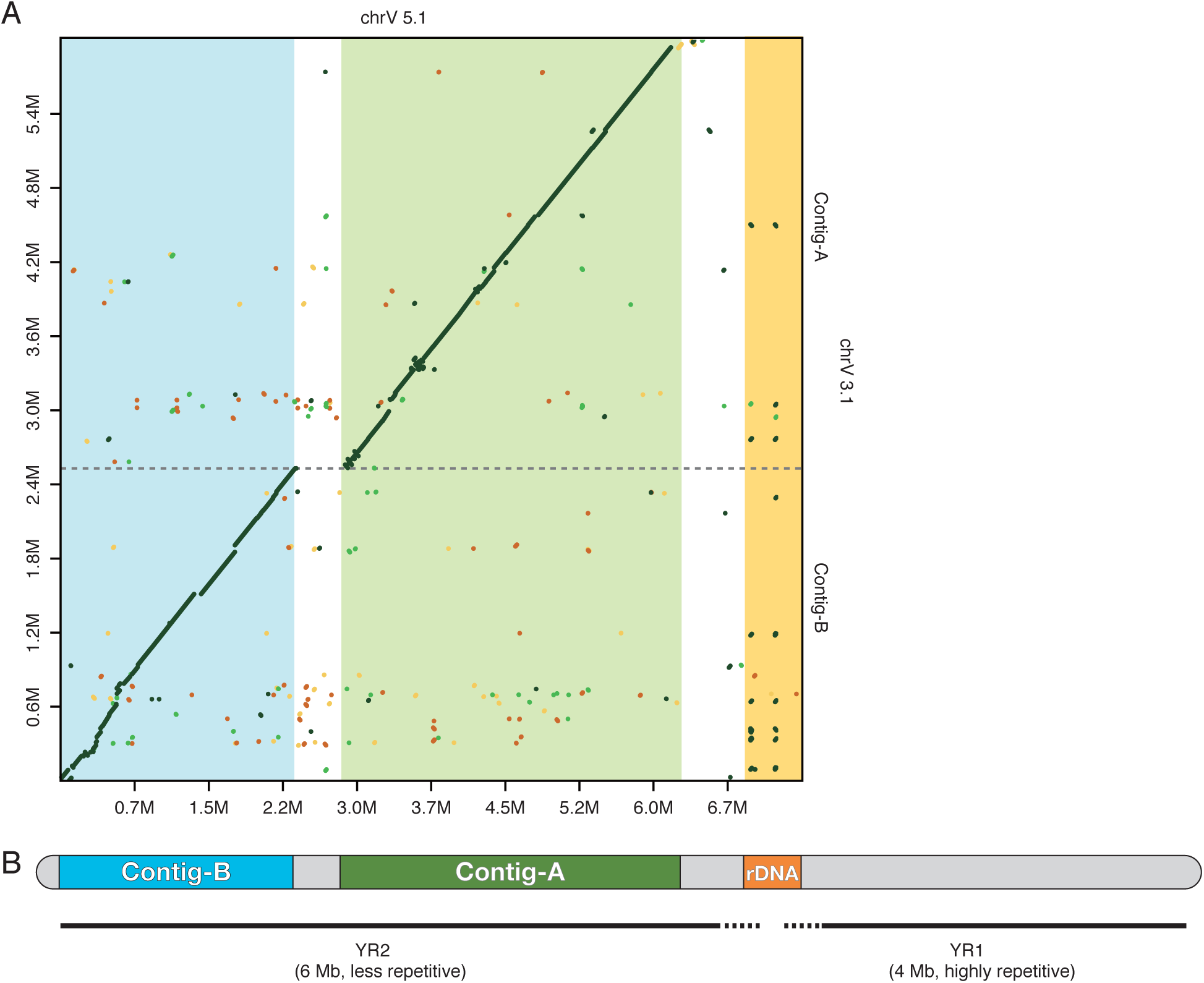
Chromosome V structure. (A) Comparison of the chromosome V sequences from the genome assembly versions 5.1 (this study) and 3.1 (Bowman et al. 2017). The regions corresponding to those previously sequenced (Contig-A and Contig-B; Yamato et al. 2017) and an rDNA cluster are colored. The rDNA cluster contains 6 copies of rDNA repeat unit, which shows 99.6% and 97.0% similarities to the autosomal and U-chromosomal copies (Fujisawa et al. 2003), respectively. (B) Schematic diagram of the V chromosome structure. The V-chromosomal segments, YR1 and YR2, identified in the previous study (Yamato et al. 2017) are represented by thick lines. Note that the boundary between YR1 and YR2 is not determined.

**Supplemental Figure 3.**
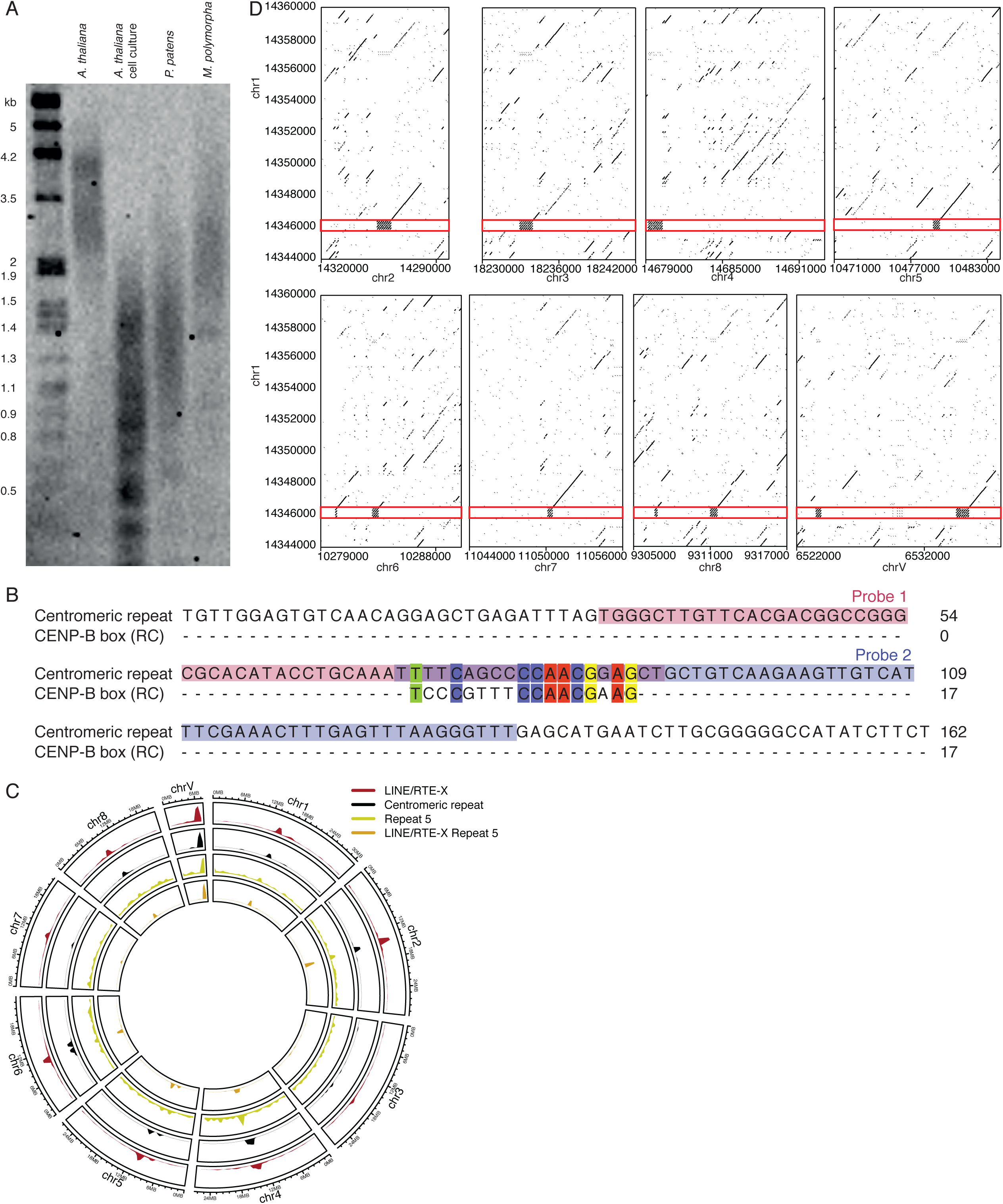
Centromeres and telomeres in *Marchantia*. (A) Comparative TRF (telomere repeat fragment assay, Southern blotting) analysis of DNA isolated from *Arabidopsis thaliana* Col-0 ecotype, *Arabidopsis thaliana* cell culture, *Physcomitrella patens* strain Gransden and *Marchantia polymorpha*. As expected, telomeres in all plants display a heterogeneous profile, but the mean length differs between the species. *M. polymorpha* telomeres (mean TRF 2,058 bp) are shorter than in *A. thaliana* (mean TRF 2,976 bp), but longer than in the model moss *P. patens* (mean TRF 1,443 bp). Telomere lengths of *A. thaliana* cell culture are shorter than in plants, as previously demonstrated [2].The blot was hybridized with a DIG-labeled (TTTAGGG)_4_ probe. Molecular weight markers are shown on the left. (B) Alignment of centromeric repeat sequence and reverse complemented CENP-B box sequence. Matching residues shown in colour. Probes used for FISH experiments are highlighted. (C) Circos plot of centromere-related feature distributions across the genome. Each band shows the density of each feature per chromosome, relative to the greatest density per band. Centromeric repeat band based on positions of BLAST hits with E-values < 10^-30^ using the putative centromeric as a query against the *Marchantia* genome. LINE/RTE-X Repeat 5 band corresponds to all LINE/RTE-X elements belonging to repeat cluster 5. (D) Dot plots between chromosome 1 and each other chromosome. Putative centromeric repeat is highlighted on chromosome 1. Sequences with a score greater than 50 over 10bp windows appear as dots. Genomic coordinates shown along the x-axis and are the same for chromosome 1 in each plot.

**Supplemental Figure 4.**
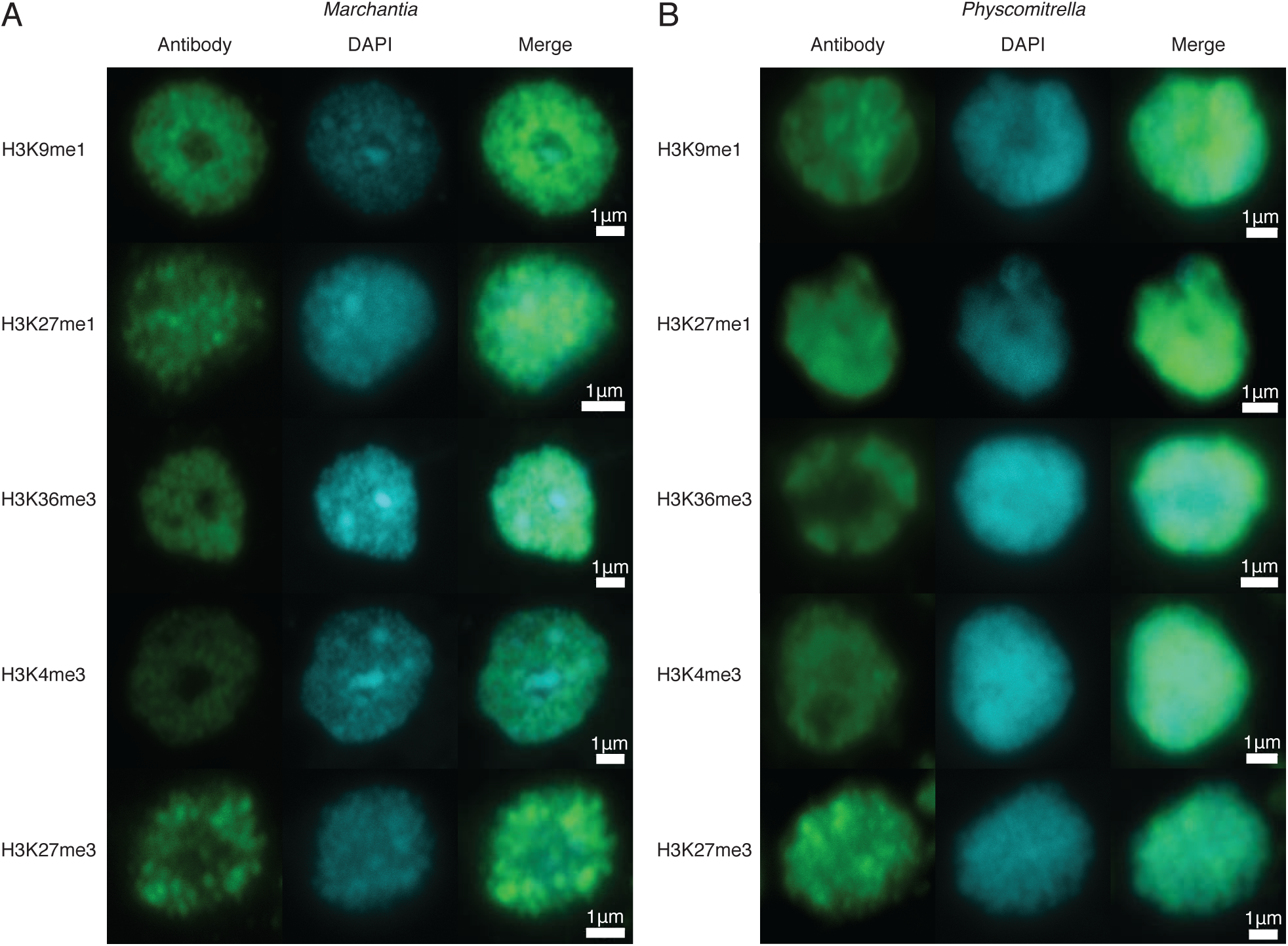
*Marchantia* and *Physcomitrella* nuclei immunostaining. (A) Immunostaining of isolated *Marchantia* nuclei. Green is the indicated chromatin mark. Blue is DAPI-stained DNA. (B) Immunostaining of isolated *Physcomitrella patens* nuclei. Green is the indicated chromatin mark. Blue is DAPI-stained DNA.

**Supplemental Figure 5.**
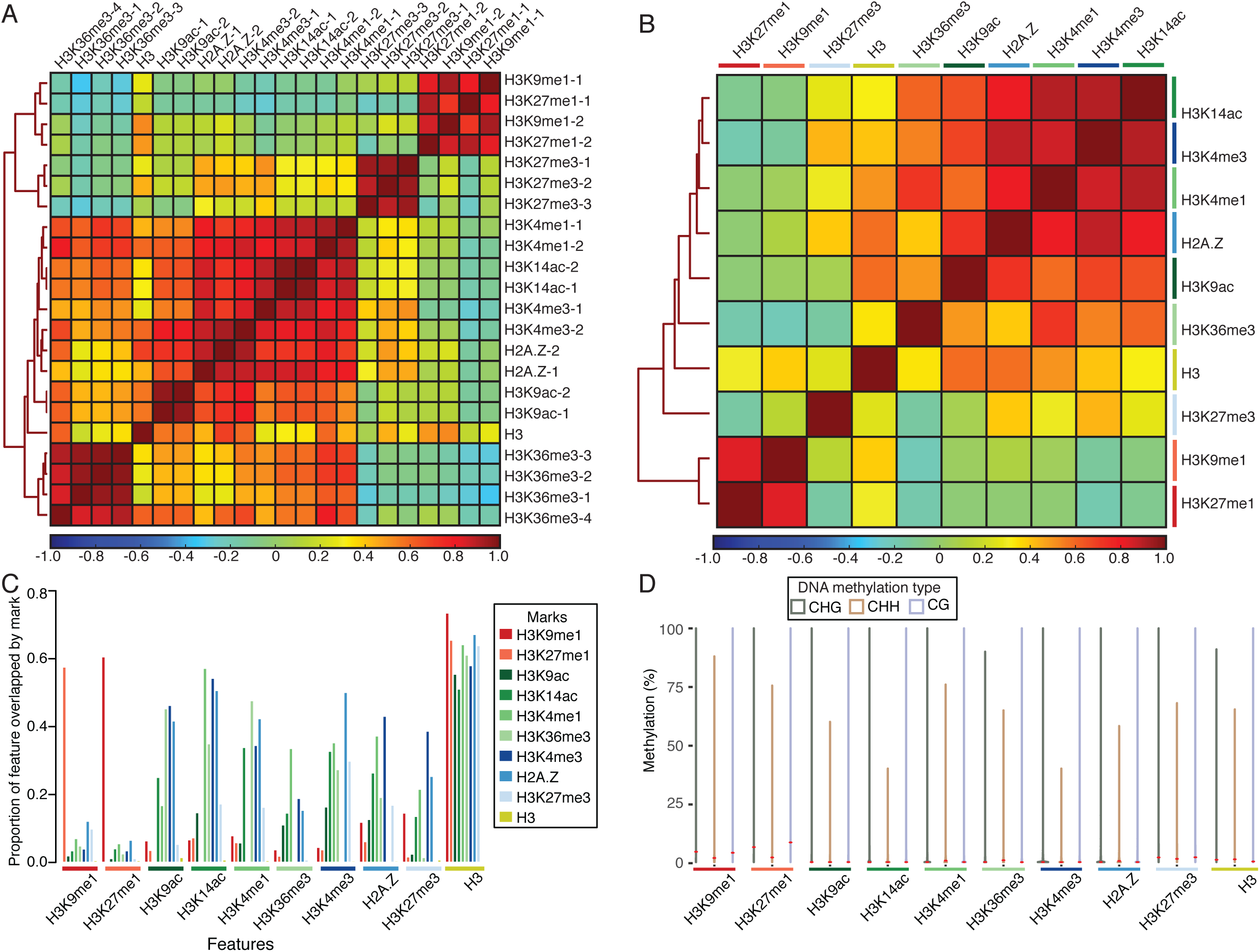
Distribution of chromatin marks in the *Marchantia* genome. (A) Pearson correlation heatmap of CUT&RUN biological replicates. Colours represent the correlation coefficient, with red for high similarity and blue for low similarity. Hierarchical clustering shown to the left of the heatmap. (B) Pearson correlation heatmap of merged CUT&RUN samples. Colours represent the correlation coefficient, with red for high similarity and blue for low similarity. Hierarchical clustering shown to the left of the heatmap. (C) Proportion of chromatin mark peaks overlapped by other chromatin mark peaks. The total length of overlapping chromatin mark peaks was divided by the total length of peaks of chromatin marks (features; along x axis) to determine the proportion of feature lengths overlapped by other chromatin marks. (D) DNA methylation levels over chromatin mark peaks. Methylation percentage calculated per chromatin mark peak. Width relative to density of peaks. Red dots indicate median methylation values.

**Supplemental Figure 6.**
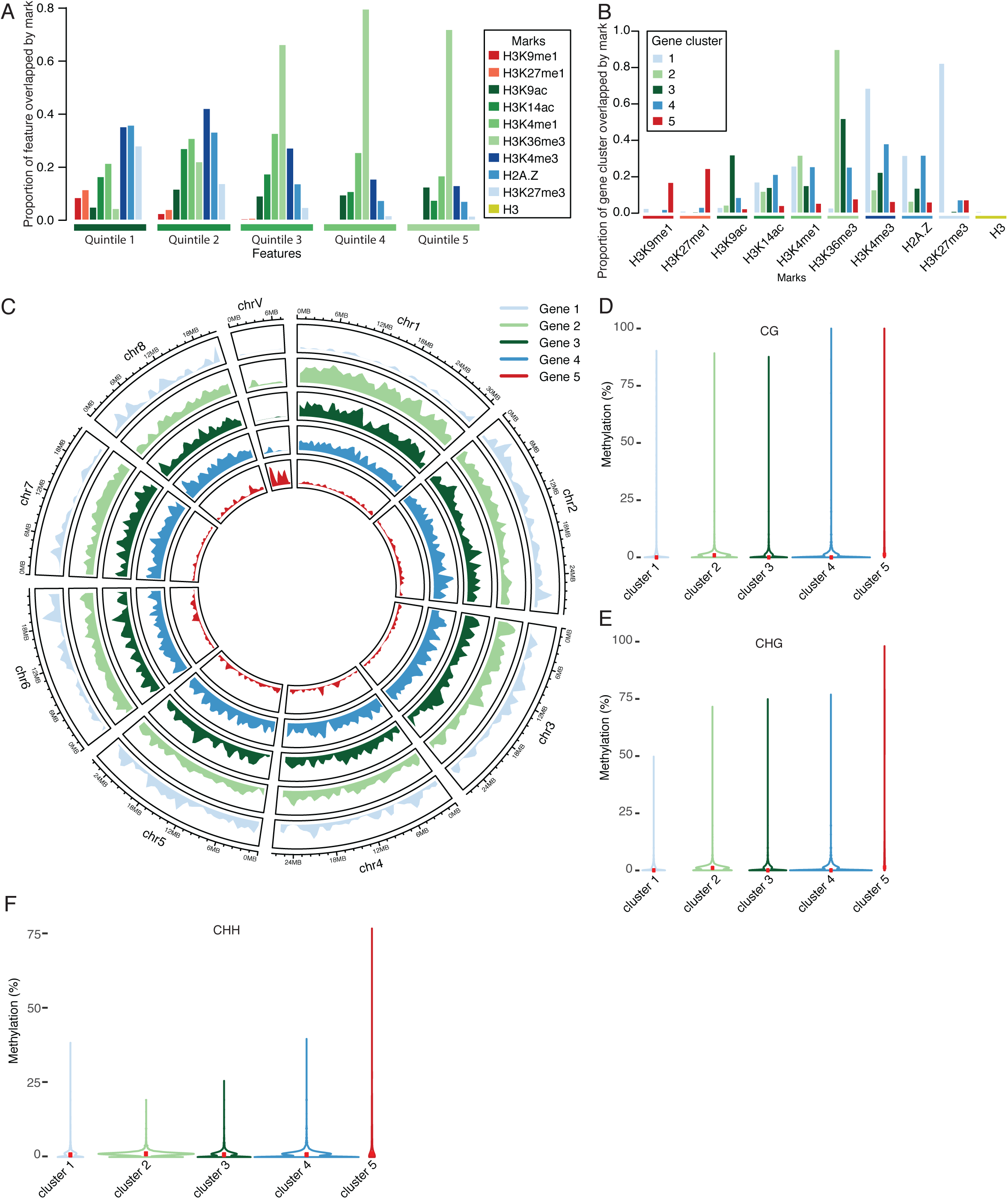
Association of chromatin marks with genes. (A) Proportion of genes per expression quintile overlapped by chromatin mark peaks. The total length of chromatin mark peaks overlapping genes was divided by the total length of genes per quintile to determine each proportion. Quintiles correspond to transcript per million values as follows: 1: 0-0.073; 2: 0.073-2.013; 3: 2.013-12.410; 4: 12.410-33.950; 5: 33.950-23567.63. (B) Proportion of gene clusters overlapped by chromatin mark peaks. The total length of chromatin mark peaks overlapping genes per gene cluster was divided by the total length of genes per gene cluster to determine each proportion. (C) Circos plot of gene cluster distribution across the genome. Each band shows the density of genes per gene cluster per chromosome, relative to the greatest density per band. (D) DNA CG methylation levels of gene clusters. Methylation percentage calculated per gene in each gene cluster. Width relative to density of genes. Red dots indicate median methylation values. (E) DNA CHG methylation levels of gene clusters. Methylation percentage calculated per gene in each gene cluster. Width relative to density of genes. Red dots indicate median methylation values. (F) DNA CHH methylation levels of gene clusters. Methylation percentage calculated per gene in each gene cluster. Width relative to density of genes. Red dots indicate median methylation values.

**Supplemental Figure 7.**
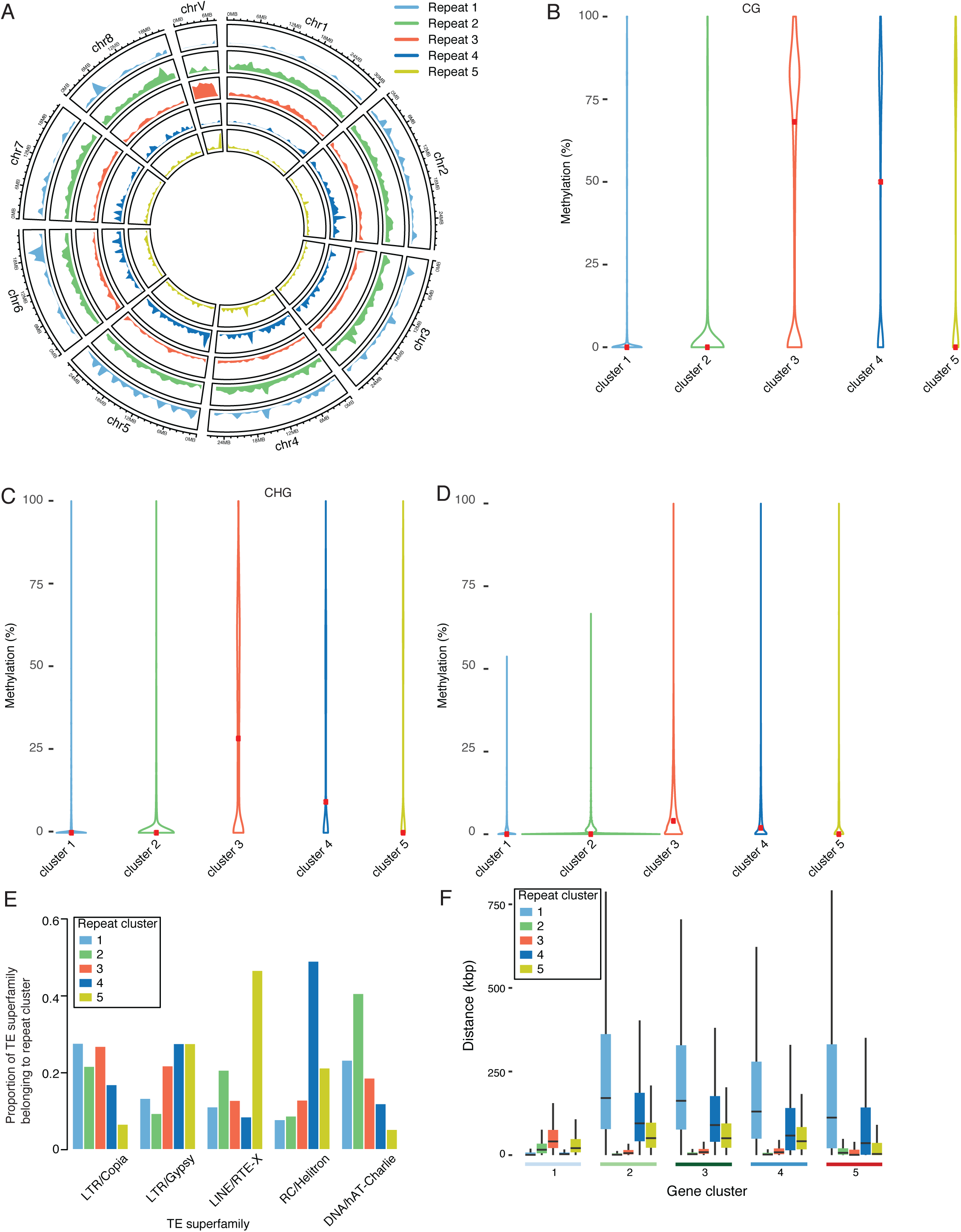
Association of chromatin marks with transposons. (A) Circos plot of repeat cluster distribution across the genome. Each band shows the density of transposons per repeat cluster per chromosome, relative to the greatest density per band. (B) DNA CG methylation levels of repeat clusters. Methylation percentage calculated per transposon in each repeat cluster. Width relative to density of transposons. Red dots indicate median methylation values. (C) DNA CHG methylation levels of repeat clusters. Methylation percentage calculated per transposon in each repeat cluster. Width relative to density of transposons. Red dots indicate median methylation values. (D) DNA CHH methylation levels of repeat clusters. Methylation percentage calculated per transposon in each repeat cluster. Width relative to density of transposons. Red dots indicate median methylation values. (E) Proportion of transposon superfamily length belonging to repeat clusters. The total length of transposon superfamilies belonging to repeat clusters was divided by the total length each repeat cluster and scaled per transposon superfamily to determine each proportion. (F) Boxplot of distances between each gene and the nearest transposon per repeat cluster. Briefly each gene is compared to all transposons belonging to a repeat cluster to find its nearest neighbor. Genes are divided based on the gene cluster they belong to. Distances in kilobases (kbp). Coloured boxes represent interquartile range and lines represent median values. Outliers not shown.

